# Information Flow and Traveling Waves show distinct cortical trajectories

**DOI:** 10.64898/2026.08.21.746280

**Authors:** Andrea Alamia, Antoine Grimaldi, Andres Canales-Johnson, Misako Komatsu, Frederic Chavane, Martin Vinck

## Abstract

Traveling waves (TWs) are a hallmark of cortical dynamics, proposed to organize neuronal activity across space and time. Yet their propagation can oppose the causal flow of information, challenging their functional interpretation. We demonstrate this using computational simulations and validate our predictions in electrophysiological recordings. We introduce the Directional Information Flow Field (DIFF), which adapts Granger causality to two-dimensional neural recordings to map directed interactions across cortical networks. In computational models, DIFF recovers the direction and sources of information flow, whereas delays and inhibition can cause TWs to propagate in opposing or orthogonal directions. We validate these findings in human and non-human primate electrophysiological recordings, revealing reversed or orthogonal flow axes, distinct spatial origins, and frequency-dependent differences in feedforward and feedback dynamics. These findings challenge the functional interpretation of TWs and establish DIFF as a complementary framework for revealing directed neural communication underlying cortical dynamics.

## Introduction

Neural activity propagates between and within brain regions as traveling waves (TW) at macroscopic and mesoscopic scales (Ermentrout and Kleinfeld, 2001; Muller et al., 2018; Ye et al., 2023), both spontaneously (Davis et al., 2020; Nauhaus et al., 2009) and during sensory (Benucci et al., 2007; Campbell et al., 2024; Muller et al., 2014; Pang et al., 2020) and motor processing (Hatsopoulos et al., 1998; Rubino et al., 2006). Increasing evidence suggests that traveling waves constitute a fundamental organizing principle of neural dynamics (Cruddas et al., 2026; Pesaran et al., 2018; Richardson et al., 2005; Xu et al., 2023), and have been implicated in diverse cognitive functions, including perception (Davis et al., 2020; Luo et al., 2021), attention (Alamia et al., 2023; Galas et al., 2025), and memory (Lubenov and Siapas, 2009; Patel et al., 2012; Zeng et al., 2024; Zhang et al., 2018). Existing approaches predominantly identify traveling waves from the spatial organization of oscillatory phase, thereby providing a quantitative description of wave propagation across multiple spatial scales (Alamia and VanRullen, 2019; Alexander et al., 2013; Das et al., 2023; Davis et al., 2020; Lefèvre and Baillet, 2009; Muller et al., 2014; Schwenk and Alamia, 2026).

Although phase-based approaches have substantially advanced the characterization of traveling waves, they do not explicitly incorporate measures of directed interactions between neural populations. Causal inference methods, including Granger causality (Granger, 1969) and Transfer Entropy (Duan et al., 2013), provide a complementary framework for quantifying directed interactions between neural populations. Here, we transform pairwise Granger causal influences into a spatial vector field, the Directional Information Flow Field (DIFF), enabling traveling waves to be characterized in terms of both their phase organization and the directionality of neural interactions. Using computational models, we show that phase-based and causal descriptions can diverge when information propagates through inhibitory pathways or networks with asymmetric delays, whereas Granger causality accurately recovers the direction of information flow in the linear systems considered here. To check if such divergence actually occurs in real experimental data, we applied DIFF to marmoset ECoG (Canales-Johnson et al., 2021) and human EEG (Pang et al., 2020) recordings. Our results reveal organizational features of cortical traveling waves that complement conventional phase-based analyses, including differences in propagation axes, wave origins, and the relationship between oscillatory frequency and feedforward or feedback processing.

## Results

To motivate the development of a Granger-based TW analysis (DIFF method), we first investigated 1D models in which an external input is introduced at one node, generating a ground-truth information flow outward from this node and propagating through either excitatory (E) or inhibitory (I) connectivity (defining four conditions, as shown in Figure 1A). Further below, we also consider 2D models (Figure 1B). We systematically compared how GC and phase-based TW analyses capture the ground-truth direction of the information flow in different cases.

**Figure 1:**
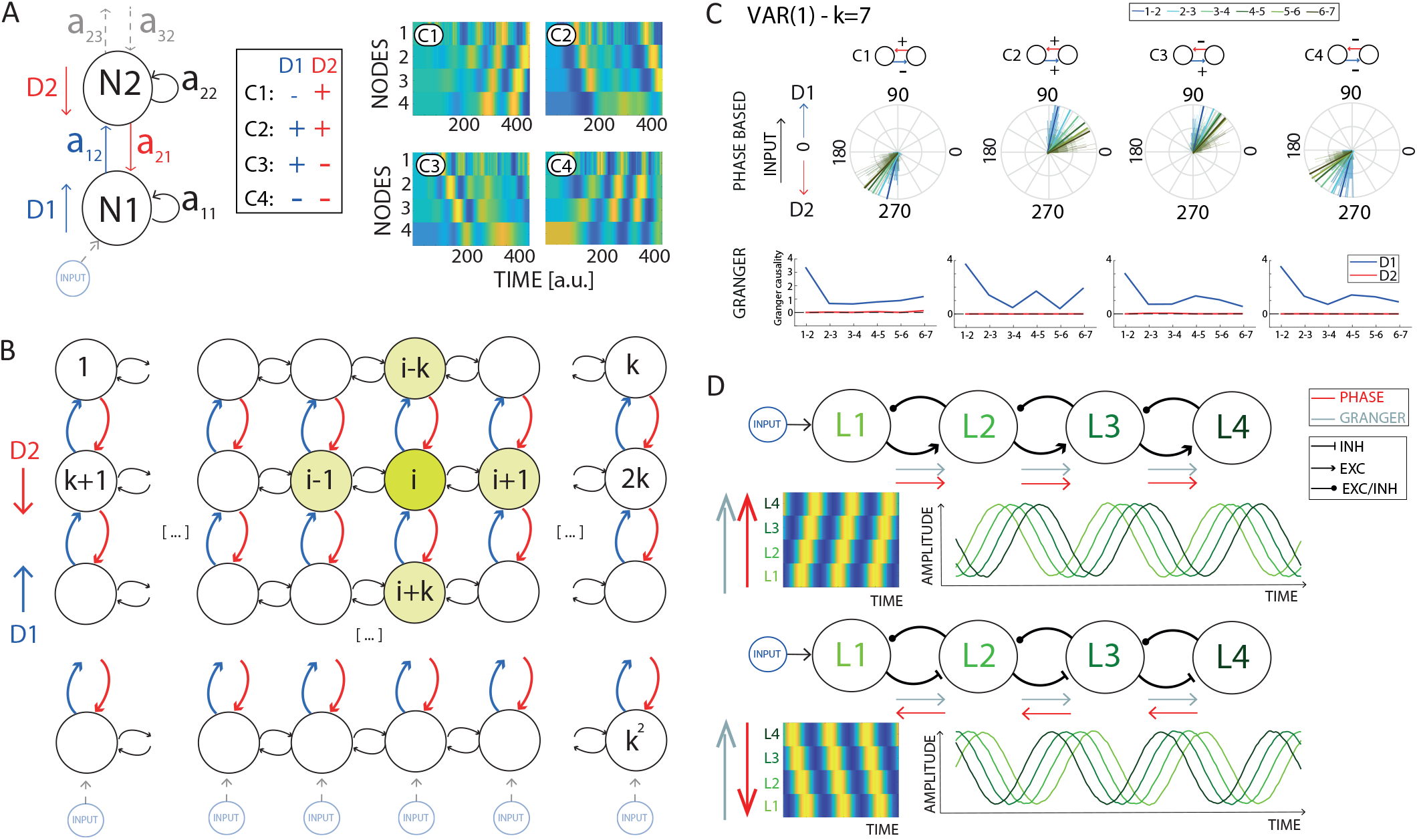
A) Schematic representation of the VAR models, in which subsequent nodes are connected via parameters *a*_*i,k*_. We considered four cases in which the connections in both directions either have positive or negative signs, as summarized in the table. We defined D1 as the direction from the perturbed node, and D2 as the opposite direction (in blue and red, respectively). The four rightmost panels show 2D representations of the activity in the four cases of the cortical model with 4 nodes. The x-axis and y-axis represent time and levels (i.e., nodes), respectively. The color (in arbitrary units) reflects the activity over time in each node. B) Schematic representation of the two-dimensional VAR model composed of *k*^2^ nodes. Each node is connected to its neighbors. A perturbation to the nodes in the lowest edge defines the direction D1 and D2 parallel to the perturbation (in red and blue in the figure). As in the 1D case, we defined four conditions by varying the signs of the D1 and D2 connections. C)Results for the VAR(1) case for phase and Granger analysis, for k=7 nodes. Each panel shows a case, as described in the upper insert, consisting of a rose plot showing the average phase difference in degrees (90^°^ indicates a flow toward D1, 270^°^ toward D2), and a plot showing pairwise Granger causality between subsequent nodes. In all cases, Granger values reveal a stronger directionality in the D1 direction, whereas phase differences reveal results opposite to those expected in cases 1 and 4. D) Summary of the simulations’ results. The upper and lower plots outline the different cases from Figure 1. When the connections in the D1 direction are inhibitory, the phase-based and Granger-based analyses yield different results.

### Phase-based for the 1-D VAR(1) case

For the 1D case shown in Figure 1A, we first analyzed the phase differences between subsequent nodes. As the first node was perturbed by the external input, we expected traveling waves to emanate in Direction 1: i.e. from the first to the last node. However, as seen from the rightmost panels in Figure 1A, the direction of phase-based TWs was highly dependent on the sign of connectivity between the nodes. In particular, phase-based TWs emanated in Direction 1 from the first (lower) node to the last (higher) node when the connections in Direction 1 were excitatory (i.e., *a*_*i,i*+1_ > 0). By contrast, TWs emanated in Direction 2 when the Direction 1 connections were inhibitory (i.e., *a*_*i,i*+1_ < 0).

To systematically quantify these differences, we computed the relative phase differences between subsequent nodes. We found a negative phase difference between all subsequent nodes when the *a*_*i,i*+1_ coefficient was negative (i.e., Cases 1 and 4), indicating that the propagation was backward from higher to lower nodes (Figure 1C). We observed an opposite direction flow when the *a*_*i,i*+1_ coefficients (Direction 1) were positive (in the second and third cases, i.e., a positive phase difference between subsequent nodes). This result was supported by a Von Mises test (V-test) for non-uniformity of circular data considering the mean direction in any pair of nodes (in all conditions and pair of nodes, considering models with k=3 and k=7 nodes, we obtained for k=3 all V-values > 250 and *p* < 0.0001; and for k=7 all V-value > 180 and *p* < 0.0001, with k being the number of layers considered). We found consistent results for k=3 and k=7 nodes (Figure 1C, and S1). All in all, the results from the phase-based analysis suggest that the direction of propagation, as measured by the phase difference between subsequent nodes, depend on the sign of the *a*_*i,i*+1_ coefficient. A positive influence from the *i* to the *i*+1 node determines the direction of phase-based propagation from lower to higher nodes (i.e., in direction D1, 1A), whereas a negative modulation produces a direction flowing in the opposite direction, from higher to lower regions (i.e., direction 2).

### Granger-causality analysis for the 1-D VAR(1) case

We then applied Granger causality analysis to assess the direction of information flow in comparison to phase-based analyses. The lower panels in Figure 1C show the GC values in the time domain between all nodes, with positive GC values indicating Granger-causal flow in Direction 1. We found highly similar patterns of GC influences for all four cases, regardless of the signs of the connections. In particular, GC analyses indicated much stronger Granger-causal influences in Direction 1, emanating from the perturbed node, as compared to Direction 2, for which Granger-causal influences were negligible. For example, for k=3, we found GC values of 2.566 ± 0.38 from node 1 to node 2 and 1.019 ± 0.06 from node 2 to node 3, while GC values in the opposite Direction 2 were smaller than 0.08. Similarly, when k=7, we found the larger GC value from node 1 to node 2 (2.880±0.31), whereas all other GC values from node *i* to node *i* + 1 (with 2 ≤ *i* ≤ 7) were between 0.925 ± 0.14 and 0.988 ± 0.15. By contrast, all the GC values connecting nodes in Direction 2 were smaller than 0.02 ± 0.02. Note that, as expected, most connections did show a significant effect when compared against a theoretical asymptotic null distribution (Barnett and Seth, 2014): This is due to the fact that a small innovation error E(*t*) (i.e., additive, Gaussian noise) was added as an input to all of the nodes.

Together, these results suggest that the Granger and the phase-based analysis provide complementary information: GC reveals the causal flow of information, following the input information propagation starting from the perturbation of Node 1, whereas the second one, by construction, provides information about the evolution of the phase, which we found to be strongly influenced by the sign of the connectivity. We generalized these observations to VAR(k) models where *k* ∈ {2, 3, 4} and with different temporal delays (see supplementary materials and Figure S1)). Furthermore, we generalized these results to a cortical model grounded in the hierarchical predictive model of inter-areal interactions (Alamia and VanRullen, 2019), which includes biologically plausible temporal delays between subsequent cortical nodes (see supplementary materials and Figure S1D). In all cases, GC analyses were consistent with the ground-truth direction of information flow, from the input layer and traveling along the subsequent nodes, regardless of the sign of the connection weights between nodes, whereas phase-based TWs were strongly modulated by the sign of the connectivity as summarized in Figure 1D.

### Asymmetric Delays

We then investigated, using VAR processes with asymmetric delays, whether the discrepancies between the phase-based and GC analyses are primarily due to the presence of inhibitory connections or can also emerge due to differences in the delays between the two opposite D1 and D2 directions. For the phase-based analysis, the upper row of Figure S2 shows that the results obtained for Case 1 and Case 3 (i.e., opposite signs between directions) were consistent for almost all pairs of delays, with the relative phase representing the averaged phase differences between subsequent nodes. The relative phase-change converged to 0 for high temporal delay. However, Cases 2 and 4 (same signs) showed an inversion of the phase gradient when the delays in D1 are much greater than the delays in D2. Even though the phase inversion was consistent for multiple pairs of delays, most relative phases for *δ*_1_ > 15 and *δ*_2_ < 15 were included in a relatively small interval between [ − 0.08π, 0.08π] (for clarity, the colorbar in Figure S2 is in logscale). Regarding the Granger analysis (bottom row), coefficients were positive and of comparable amplitude independent of the configuration, confirming an overall flow of information following the direction of the perturbation D1. Altogether, these results corroborate and extend the conclusion that the Granger and the phase-based analysis provide complementary information about the flow directionality when the connections in the D1 direction are inhibitory and when strong asymmetric delays are present between feedforward and feedback connectivity.

### 2D networks and DIFF method

We then analyzed cases in which the nodes are spatially organized in 2D networks and locally connected to their neighbors, as shown in Figure 1B. To this end, we developed the DIFF method to characterize TWs based on Granger-causality, by (1) computing Granger-causality values from and toward a node and its neighbors, (2) computing the spatial resultant vector from each node, and (3) computing a vector field across all nodes. We refer to this vector field as the Directional Information Flow Field (DIFF, see methods). The Divergence, a scalar that quantifies the gradient of this vector field across *x* and *y* dimensions, can then be used to find sources of TWs in a 2-D array.

In our simulations, we considered two distinct conditions in which we continuously perturbed two subsets of nodes: in the first, the external perturbations affected two non-adjacent nodes (Fig. 2); in the second, all the nodes on the lower edge were perturbed (Fig. S3). We computed the phase vector field and the Directional Information Flow Field (DIFF, see methods) in both conditions for all four cases, and we computed the absolute divergence in both vector fields to identify the source of the external perturbations. As shown in Figure 2A, the phase analysis provided qualitatively different vector fields in the four cases. In Cases 1 and 3, where the connections had opposite signs, the phases revealed a flow in the excitatory direction running through the 2 input nodes (D2 and D1 in Cases 1 and 3, respectively). In Cases 2 and 4, in which all connections share the same sign, the phases show either an outgoing pattern from the sources in the case of excitatory connections (Case 2) or a more random pattern if all connections are inhibitory (Case 4). To statistically analyze the results, we tested whether the divergence value at the perturbed node was significantly different from that in all other nodes (Figure 2C). We consider all divergence values as a univariate Gaussian distribution and computed the probability (i.e., the p-value) that the divergence values at the perturbed nodes are sampled from the same distribution. We found significant p-values *p* < 0.05 only for case 2, whereas all other p-values were larger than 0.05. This result was consistent and robust to an increase in the amount of independent Gaussian noise E introduced in all nodes, as shown in the upper panels of Figure 2C. This analysis suggests that, in a 2D network, the divergence of the phase-based vector field can reveal the source of propagating activity if all connections are excitatory (Case 2), but not otherwise.

**Figure 2:**
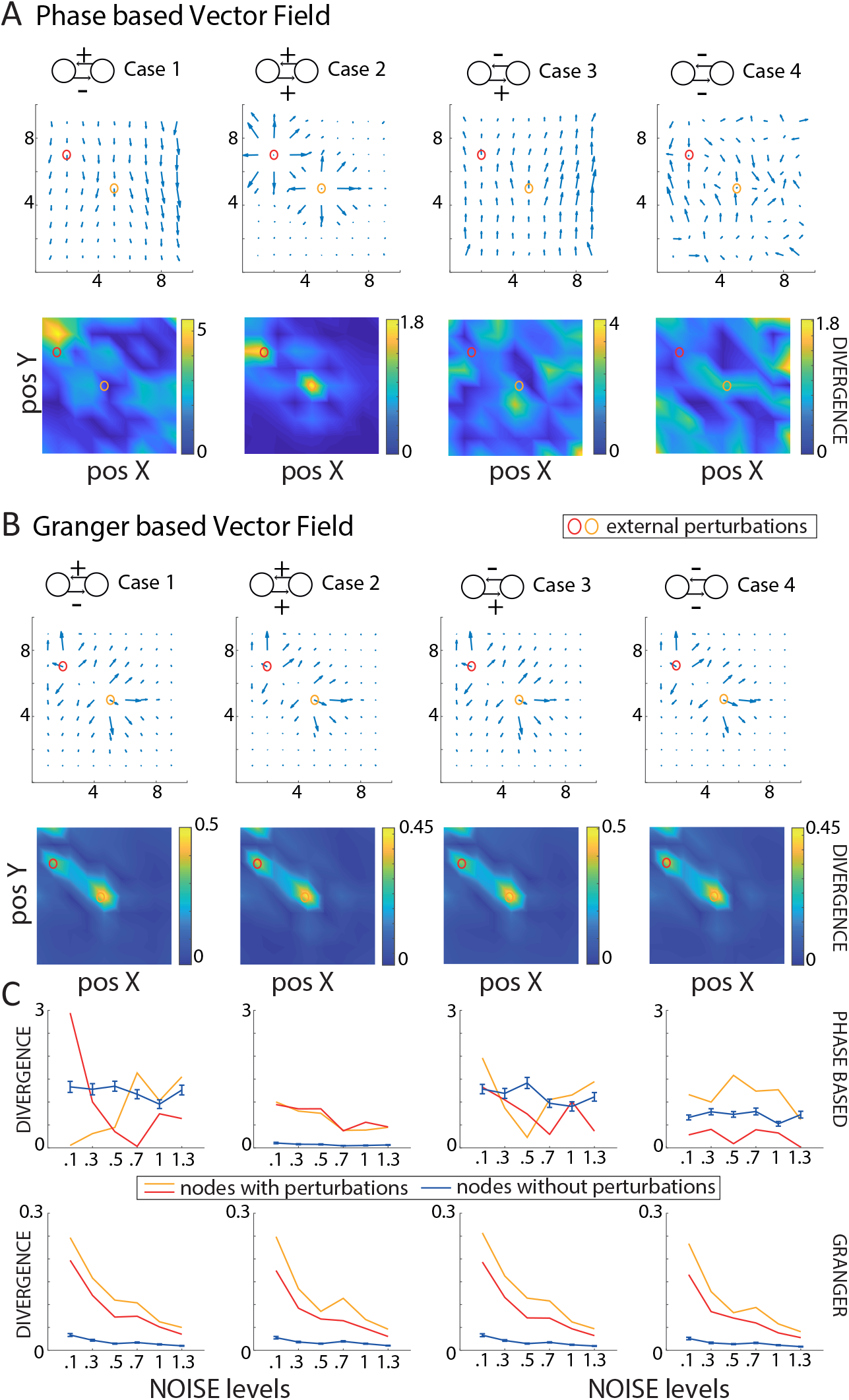
Results for the 2D networks. Panels A and B show the results of the phase-based and Granger-based (i.e., DIFF) vector fields. Each column corresponds to one of the four cases, as shown in the inserts above. In each panel, the first and second rows represent the vector field and its divergence, respectively. The two nodes circled in red and yellow are the sources of the perturbations. Panel C) shows the divergence as a function of the noise *ϵ* introduced independently in all nodes. Blue lines represent the divergence at the nodes without perturbation (mean ± standard deviations), whereas the colored lines represent the divergence in the perturbed nodes. Only the Granger-based DIFF method identifies nodes with perturbations in all conditions, and robustly to noise.

In contrast to the phase analysis, the Directional Information Flow Field (DIFF) yielded consistent results across cases (Figure 2B), indicating an outward pattern from the external perturbation. In particular, the divergence values at the perturbed nodes were significantly different from the divergence values at all the other nodes for all cases (all *p* < 0.001). These results were consistent with the increase of the noise levels E introduced in each node. Even though the divergence values at the perturbed nodes decreased with the noise level (Figure 2C), the difference between perturbed nodes and all the other ones remains significant (all *p* < 0.01). We then confirmed and extended the previous results to the case of a planar wave propagating from the lower edge of the network (akin to Node 1 in the 1D case), as shown in Figure S3. In particular, we observed that the DIFF method applied to the Granger-based vector field reliably identified the perturbed nodes in all cases, as indicated in Figure S3B, and irrespective of the amount of noise level E (S3C). By contrast, the phase-based vector field indicated the correct direction in Cases 2 and 3 (as in the previous 1D analysis), but an opposite pattern in Cases 1 and 4 (specifically closer to the perturbed node), similar to the 1D model. Different from the Granger-based vector field, the divergence of the phase vector field did not identify the source of the planar wave. In summary, across all simulated conditions - 1D and 2D architectures with different delays - phase-based and Granger-based TW analyses yielded dissociable and complementary results, with only the latter consistently and reliably identifying the true sources of perturbation across conditions. In the following, we compare the two methods in a range of different experimental recordings.

In what follows, we apply these methods to datasets from marmosets and humans to study differences between phase-based methods and DIFF and to uncover directional TWs using the DIFF method. We show that prominent differences between phase-based and Granger causality methods can arise in real data and are not merely artifacts of overly constrained or unrealistic theoretical assumptions.

### Comparison between phase-based and Granger-based TWs in human EEG data

We first analyzed a scalp EEG dataset recorded from 13 human participants performing a visual task in which five-second periods of visual stimulation alternated with a blank screen (Pang et al., 2020). Previous analyses of traveling waves reported alpha-band [8-12 Hz] activity propagating from frontal to occipital regions during rest (Alamia and VanRullen, 2019; Mohan et al., 2024; Park et al., 2025), with bidirectional propagation (from occipital to frontal and vice versa) during visual stimulation (Alamia et al., 2023; Pang et al., 2020). The phase-based vector field analysis corroborates these earlier findings. The upper panels of Figure 3A show the across-participant average phase vector field during the rest phase of the task, in which participants kept their eyes open but without any visual stimulation presented. The vector field showed a top-down propagation for alpha- and beta-band oscillations (between 8-12 Hz and 13-30 Hz, respectively), as indicated by negative divergence values in central regions. By contrast, the phase vector field indicated a bottom-up propagation pattern for gamma oscillations (above 30 Hz, hence broadband (BB) rather than narrow-band gamma), as well as a spiral propagation pattern in occipital regions in the theta band. In contrast, the DIFF method revealed a distinct medio-to-lateral propagation pattern in frontal and central regions across the alpha, beta, and gamma bands, with positive divergence values indicating a central source around the midline. Vector fields derived using the DIFF method exhibited substantially greater similarity across frequency bands than phase-based vector fields (Figure S9A). During both visual stimulation and rest (stimulus ON and OFF conditions), cosine similarity values for the DIFF method remained consistently elevated (mean ~ 0.6), whereas phase-based vector fields showed markedly lower similarity across bands (generally ≤ 0.2). These findings indicate that the DIFF approach captures dynamics that are more conserved across frequencies, consistent with a frequency-independent process, whereas phase-based methods reveal dynamics that are more strongly frequency-specific. This distinction was further supported by the absence of correlation between divergence measures obtained with the two approaches, together with the relatively homogeneous divergence values across bands for the DIFF method and the more heterogeneous band-dependent patterns observed for the phase-based method (Figure S9B). Figure S6 shows the average cosine similarity of vector fields across participants, revealing consistently positive values and low inter-participant variability for both the Granger-based and phase-based approaches. This was confirmed by Bayesian t-tests, providing strong evidence for values larger than 0 in all conditions (for all frequency bands: phase stimulus ON, all

**Figure 3:**
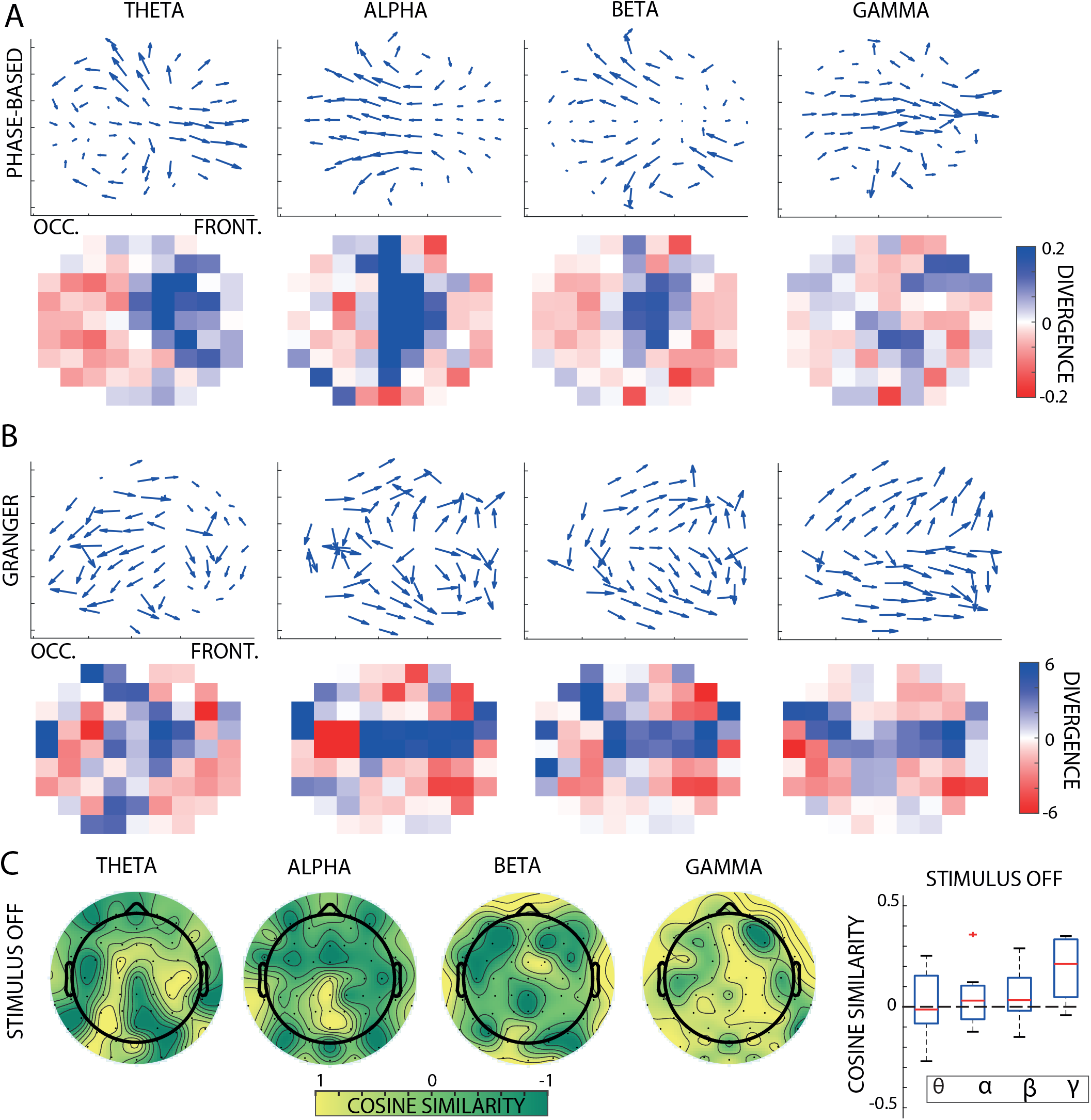
Applying phase-based and DIFF on EEG data (Pang et al., 2020). Results for scalp EEG data recorded in the absence of visual stimulation. A,B) The first row displays the between-subject average vector fields derived using the DIFF (Granger) and phase-based methods applied to scalp EEG data with visual stimulation turned off, respectively. The second row shows the corresponding average divergence of the vector fields, computed across participants. C) The plots show the cosine similarity between the DIFF and phase-based vector fields across frequency bands. With the exception of the gamma band, the two methods exhibit relatively low similarity in occipital and parietal regions. The plot on the right shows the cosine similarity averaged across electrodes for each frequency band.

*BF*_10_ > 10^5^, mean values µ ∈ [0.12, 0.29]; phase stimulus OFF, all *BF*_10_ > 10^6^; mean values µ ∈ [0.20, 0.53]; DIFF stimulus ON, all *BF*_10_ > 10^5^, mean values µ ∈ [0.12, 0.29]; DIFF stimulus OFF, all *BF*_10_ > 10^11^; mean values µ ∈ [0.23, 0.46]). We further computed cosine similarity between the DIFF and phase-based methods (Figure 3C), revealing distinct vector fields in the theta (4-7 Hz), alpha (8-12 Hz), and beta (13-30 Hz) bands, but higher similarity in the gamma-band (similar results were observed during visual stimulation; Figure S7)C. The topo-graphic plots in Figure 3C reveal a certain heterogeneity, with similarity values ranging from opposite alignment to alignment. At a global level, Bayesian t-tests across all electrodes show no difference from zero in the theta and alpha bands during and without visual stimulation (all *BF*_10_ < 0.25), and inconclusive results for the beta band (all *BF*_10_ ∈ [0.83, 1.07]). In contrast, gamma-band oscillations exhibit similar vector field patterns across the two methods, as confirmed by Bayesian t-tests against zero in both stimulus ON and stimulus OFF conditions (both *BF*_10_ > 30). We then compared the two Granger- and phase-based methods by quantifying the global directions of propagation along the fronto-occipital and medial-lateral axes for each hemisphere separately (Figure S8). Specifically, we computed the average of the real and imaginary components of all vectors in each hemisphere. Concerning the fronto-occipital axis, the Bayesian ANOVA supported an effect of stimulation (ON vs OFF) in the phase-based analysis for all frequency bands (all *BF*_10_ > 25, for theta-band *BF*_10_ = 4.78), but no effect of the hemisphere (all *BF*_10_ ∈ [0.27, 0.43]). On the contrary, the DIFF approach did not find an effect between stimulation conditions (*BF*_10_ ∈ [0.25, 0.59]), except moderate evidence in the theta-band (*BF*_10_ = 6.46). We did not report any difference between the two hemispheres (all *BF*_10_ < 0.5), and inconclusive evidence for the beta-band (*BF*_10_ = 2.12).

We found substantial differences between the two Granger- and phase-based methods along the medial-lateral axis, as shown in Figure 3 and in Figure ??. The DIFF method reveals large evidence for propagation from medial to lateral areas in all frequency bands (all *BF*_10_ > 10^6^), and no effect of the visual stimulation conditions (all *BF*_10_ ∈ [0.22, 0.64]). As illustrated in Figure 3, the DIFF approach reveals a previously uncharacterized medial-to-lateral axis of causal propagation, with a pronounced effect in frontal regions. These results demonstrate the DIFF’s ability to uncover a distinct pattern of directional information flow from scalp EEG recordings. This additional organizational dimension provides a framework for reinterpreting prior findings within a more comprehensive view of large-scale cortical dynamics.

### Comparison between phase-based and Granger-based TWs in marmoset ECoG data

Next, we analyzed a dataset consisting of ECoG recordings from electrode arrays implanted in the left or right hemispheres of one marmoset performing an auditory oddball task (Canales-Johnson et al., 2021; Gelens et al., 2024). Each trial included a 400-ms period before and a 400-ms period after stimulus onset, during which either a standard or a deviant stimulus was presented. Given the short duration of the recordings and to ensure a consistent number of cycles, we applied the DIFF and phase-based approaches to activity from the alpha to gamma bands. Note that the ECoG time series contained a spectral peak at 12 Hz (Figure S4B), and that “gamma” refers to broadband (BB) rather than narrow-band gamma, similarly to the EEG analysis. We computed vector fields for both phase-based and DIFF methods following stimulus onset. Cosine similarity between standard and deviant auditory stimuli was consistently high for both Granger- and phase-based analyses (mean across electrodes: 0.95 and 0.93, respectively). Given the strong similarity of the vector fields in all conditions, all subsequent analyses were conducted on averages.

Figure 4A shows the vector fields and their divergence in three frequency bands computed from the marmoset ECoG recordings (see Methods). To quantify the agreement between phase-based and Granger-based TWs, we computed the cosine similarity computed over the vector fields (see Methods).

**Figure 4:**
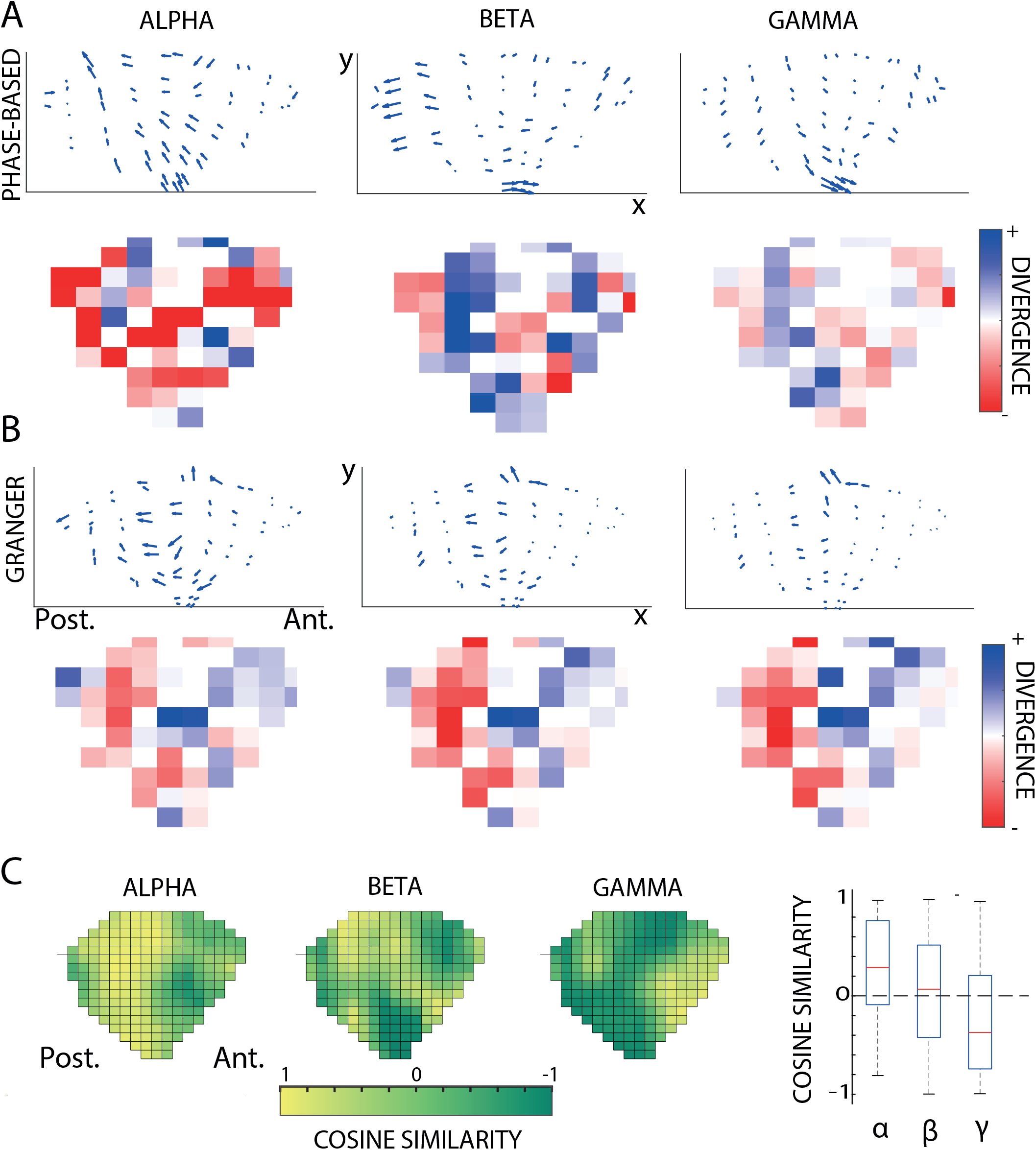
DIFF on ECoG data in Marmoset (Canales-Johnson et al., 2021; Gelens et al., 2024). A) The figure presents the outcomes of both the DIFF and phase-based methods applied to the data of one marmoset. Arrows indicate the dominant flow structure of neural interactions within each cluster. In each panel, the first and third rows depict the DIFF (Granger) and phase-based vector fields averaged across standard and deviant stimuli. The second and fourth rows display the divergence computed for each vector field. B) Topographic representation of the cosine similarity between the two methods across frequency bands. The plots indicate agreement between the DIFF and phase-based approaches in auditory areas, for alpha and beta bands, but not for gamma. Labels: Ant: anterior, Post: posterior. The rightmost boxplot shows the cosine similarity between the DIFF and the phase-based vector fields averaged across all sensors for all frequency bands. Overall, values near zero indicate that the two methods are poorly aligned.

The alpha-band (8-12 Hz) showed relatively good agreement. There were major differences across space and frequency between Granger-causality and phase-based wave patterns for all three frequency bands. The DIFF method suggests a similar TW profile for alpha, beta, and BB gamma, with the strongest flow in the anterior-to-posterior direction over the parietal cortex. This was corroborated by high cosine similarity values between frequency bands, as shown in Figure S9C. By contrast, the phase-based TW profiles differ substantially between frequency bands and from the DIFF patterns (Figure S9C.). The DIFF method revealed a propagation from more anterior regions toward occipital areas in the alpha and beta bands, as indicated by the divergence values, and less so in gamma. The phase-based approach yielded a pattern similar to that in the alpha band, as supported by cosine similarity measures between the two vector fields (Figure 4B). However, the beta-band in the phase-based vector field had a more posterior onset compared to the DIFF method, as indicated by positive divergence values in parietal and occipital regions, and a lower cosine similarity between the two methods. Gamma-band oscillations yielded distinct vector fields across the two methods, with lower overall amplitudes and divergent phase directions in the phase-based analysis. As shown in Figure 4B, the average cosine similarity across all sensors showed overall little alignment between the two methods over the whole field of view (but with large variability across electrodes), as indicated by values around zero (Bayes Factor to test difference from zeros for each frequency band α : *BF*_10_ = 0.57; β : *BF*_10_ = 0.47; γ : *BF*_10_ = 0.46, all errors <0.04%).

### Bimodal Phase Distribution of Trials Without Changes in Granger Causality

Building on our simulation results, we next asked whether the dissociation between phase-based and Granger-based vector fields held true also across trials. That is, whether variations in phase-based vector fields across trials were mirrored by comparable variations in Granger-based vector fields. Our prediction is that, if these measures are independent and complementary, as suggested by our simulations, inter-trial variability in one should not affect the other. To this end, we partitioned trials from both the Marmoset and scalp-EEG datasets based on their traveling-wave properties, as defined by phase-based analyses. In the marmoset dataset, trials were clustered using HDBSCAN on the extracted wave patterns, yielding 26.3 % and 14.4 % trials in the main two groups (3.9 % in a third group, and 55.5 % were classified as noise). In the human EEG dataset, trials were instead separated based on the direction of alpha-band propagation (bottom-up vs. top-down), a distinction widely reported in the literature (Alamia and VanRullen, 2019; Mohan et al., 2024; Park et al., 2025). We clustered trials in the stimulus ON and OFF conditions based on the mean phase direction across sensors, yielding 16.6% and 10.02% of trials with mean phase near 0 ± 0.3 radians, and 10.1% and 28.5% with mean phase near π ± 0.3 radians. Across both datasets, this stratification yielded by construction subsets of trials with clearly distinct phase-defined propagation patterns (Figure 5). We then computed Granger-based vector fields separately within each subset to assess whether differences in phase dynamics were paralleled by changes in directed functional interactions. Strikingly, the Granger-derived vector fields remained highly consistent across trial subsets, despite the pronounced differences in phase-based organization. In both marmoset and human EEG data, the overall structure and directionality of the Granger fields were largely preserved, with minimal sensitivity to phase-defined clustering or propagation direction. This was supported by the cosine similarity analysis between vector fields in the two trial subsets, as shown in Figure 5. These findings indicate that Granger-based measures capture aspects of directional information flow that are largely independent of the instantaneous phase structure underlying traveling waves. This dissociation closely mirrors our simulation results, supporting the interpretation that phase-based and Granger-based approaches complement each other, while probing fundamentally distinct dynamical features of large-scale neural activity.

**Figure 5:**
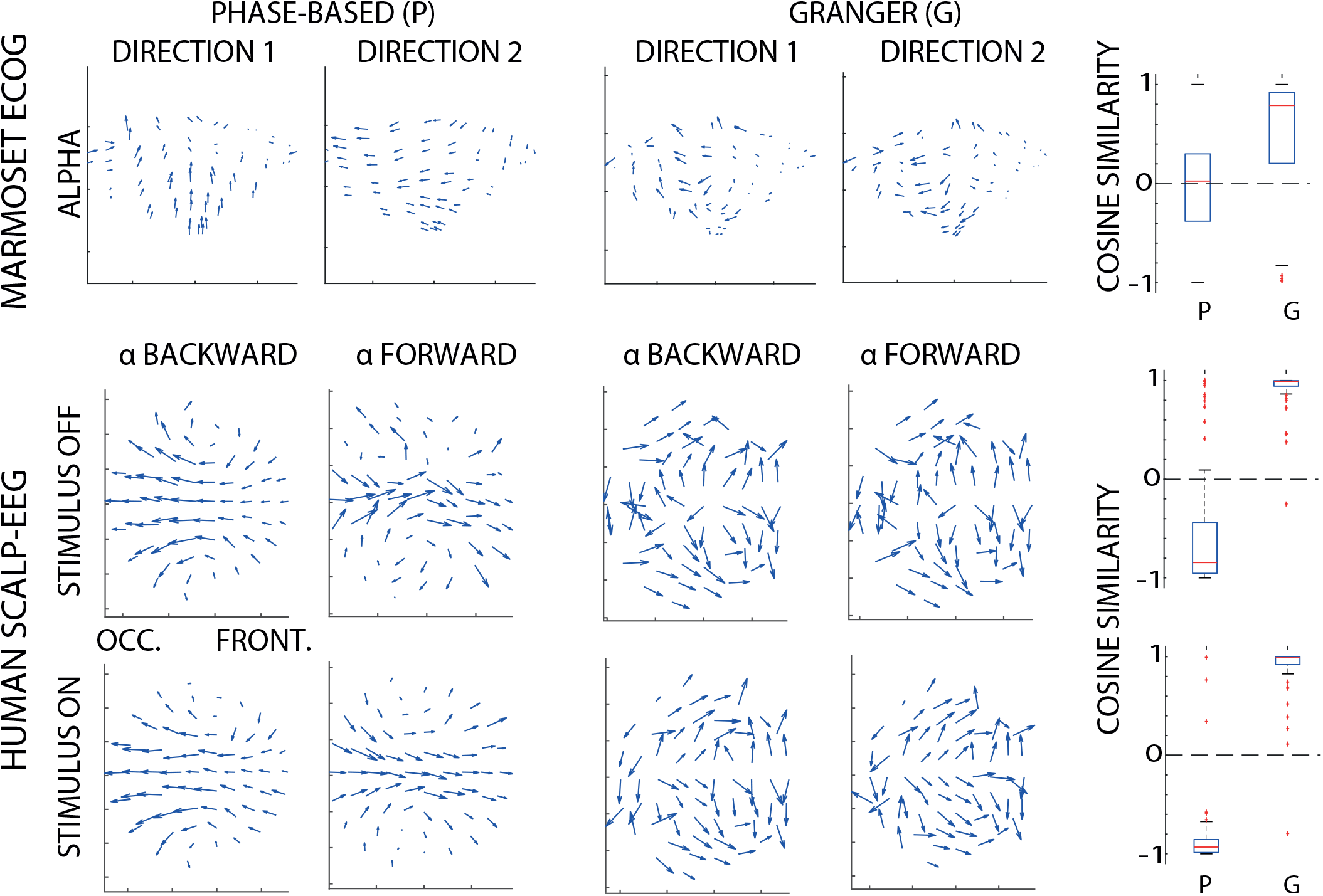
Comparison of Granger and Phase-Based Vector Fields Following Phase-Based Trial-Wise Clustering. Vector fields for the alpha band are shown for the marmoset and the human EEG data, computed using the phase-based (P) and Granger (G) methods. Regarding the marmoset data, trials are separated by directionality (Direction 1 and Direction 2) following HDBSCAN clustering on the traveling waves (i.e., phase-based) results. The panels on the right summarize the cosine similarity between vector fields computed from subsets of trials within each frequency band, shown separately for phase-based (P) and Granger (G) methods. Regarding the scalp EEG data, vector fields are computed analogously using phase-based (P) and Granger (G) approaches, organized by stimulus condition (stimulus OFF vs. stimulus ON). Within each condition, trials are split into subsets based on the forward and backward main direction of propagation. The rightmost panels show cosine similarity measures between vector fields derived from subsets of trials for each method, illustrating high consistency in the Granger case, and opposite results in the phase-based approach.

## Discussion

### Summary of the results

Spatio-temporal organization of cortical activity has been largely studied through spatial phase gradients, which do not capture the causal flow of information within the network. The relationship between TW propagation and directed information flow, therefore, remains poorly understood. Here, we show that TW propagation can dissociate from the underlying flow of information, revealing a fundamental limitation of interpreting TW direction as a proxy for neural communication. To uncover this dissociation, we estimate the directed information flow derived from Granger causality using the Directional Information Flow Field (DIFF). Using computational models, we show that phase-based and causal descriptions capture complementary aspects of wave propagation. In particular, conditions involving inhibitory interactions or asymmetric delays can dissociate the direction of wave propagation from the direction of information flow, whereas Granger causality accurately recovered the latter in the linear systems examined here. Building on these observations, DIFF transforms pairwise Granger causal influences into a spatial vector field, enabling the directionality of neural interactions to be analyzed alongside the spatiotemporal organization of traveling waves. In two-dimensional simulations, DIFF reliably localized perturbation sources. Importantly, by applying DIFF to marmoset ECoG and human EEG we revealed in real data some key organizational features of cortical traveling waves that complement conventional phase-based analyses. As predicted theoretically, we show experimentally that TW and DIFF can differ in propagation axes, spatial origins, and frequency-dependent patterns of information flow. Notably, the vector field derived from our analysis is reminiscent of previously reported spatial gradients of propagation in the brain based on gradient flows of cortical activity (Lefèvre and Baillet, 2009; Liu et al., 2026; Townsend and Gong, 2018).

### Biological plausibility of our results

Our results reveal that, under certain conditions, a significant discrepancy can arise between the identified direction of propagation of the activity, as inferred from phase differences, and the underlying causal interactions. The dissociation between phase-based and causal descriptions of wave propagation was robust across all computational models examined, including implementations with temporal delays and a biologically grounded cortical model (Alamia and VanRullen, 2019). Although our theoretical analysis focused on linear systems with homogeneous connectivity, the underlying mechanism responsible for this dissociation is not restricted to a particular model implementation. Instead, it arises when the direction of information transfer differs from the direction of phase propagation, for example, through inhibitory interactions or asymmetric transmission delays. Whether similar effects extend to strongly nonlinear cortical dynamics remains an important question, given that cortical dynamics are often modeled as nonlinear systems (Freeman and Skarda, 1985; Vinck et al., 2023). In this context, information-theoretic measures such as Transfer Entropy provide a natural extension of the present framework, enabling the estimation of directed interactions beyond the linear regime. Comparing phase-based traveling-wave analyses with Transfer Entropy, therefore, represents an important direction for future work.

At the same time, linear approximations remain highly relevant for the analysis of large-scale neural recordings. Inter-areal interactions measured with electrophysiological signals are often dominated by approximately linear components arising from afferent synaptic activity (Dowdall et al., 2023; Dowdall and Vinck, 2023; Pesaran et al., 2018; Schneider et al., 2021; Vinck et al., 2023), and local oscillatory dynamics are frequently well described as noise-driven damped oscillators (Spyropoulos et al., 2022). The assumptions adopted here may therefore capture an important component of cortical population dynamics.

Our results further highlight the potential importance of heterogeneous effective interactions at mesoscopic and macroscopic scales. Although both feedforward and feed-back corticocortical projections are glutamatergic, they recruit distinct combinations of excitatory and inhibitory neuronal populations (Callaway, 2004; Shao and Burkhalter, 1996). In particular, converging evidence suggests that feedback pathways can exert a net suppressive influence on lower cortical areas (Briggs, 2020; Isaacson and Scanziani, 2011; Murphy et al., 1999; Olsen et al., 2012; Vezoli et al., 2020; Wilmes and Clopath, 2019). Our simulations suggest that, under these conditions, top-down signals originating in higher cortical regions—for example, during imagery or predictive processing (Rao and Ballard, 1999; Rauss et al., 2011)—could propagate through feedback pathways while generating phase gradients that appear to travel in the opposite, feedforward direction. This provides a plausible physiological mechanism by which phase-based and causal descriptions of traveling waves may diverge.

Furthermore, paradoxical effects of neural activation can arise from balanced E/I networks (Tsodyks et al., 1997), as well demonstrated by inhibitory-stabilized networks (Ozeki et al., 2009), in which long-range excitation can produce net suppression of downstream activity (Chemla et al., 2019; Reynaud et al., 2012). Consistent with this view, high-frequency inputs have been suggested to predominantly target fast-spiking inhibitory interneurons through feedforward pathways (Reynaud et al., 2012; Schneider et al., 2023; Spyropoulos et al., 2024). These observations suggest that, under specific physiological conditions, the direction of information transfer may become dissociated from the direction of phase propagation. Such a dissociation may account for previous reports of mismatches between phase-based and Granger-causal measures in the beta-frequency range(Brovelli et al., 2004a), a rhythm that has been associated with inhibitory processing and top-down signaling (Bastos et al., 2020). Consistent with this interpretation, our analyses of both marmoset ECoG and human EEG revealed systematic divergences between phase-based traveling waves and DIFF, including reversed or orthogonal propagation axes and distinct spatial origins of wave activity. Together, these findings suggest that such discrepancies are not isolated observations but may instead reflect a general property of cortical dynamics when effective inhibitory interactions shape large-scale communication.

### Comparison between phase-based and Granger-based TWs in electrophysiological data

Applying DIFF to marmoset ECoG and human EEG recordings revealed several organizational features of cortical traveling waves that were not apparent from phase-based analyses alone.

Across species and frequency bands, the similarity between phase-based and DIFF vector fields was generally low, indicating that the spatial organization of oscillatory phase and the directionality of information flow need not coincide. In particular, DIFF consistently identified a dominant medio-lateral axis of directed interactions, whereas phase-based analyses predominantly emphasized anterior–posterior propagation. Moreover, substantial trial-to-trial changes in phase-based traveling waves were often accompanied by remarkably stable DIFF patterns, suggesting that large variations in phase organization can coexist with conserved patterns of directed communication. Altogether, our findings suggest that the spatial organization of oscillatory phase and the directionality of information flow can diverge, indicating that phase-based and causal analyses provide complementary views of cortical traveling waves.

The origin of these dissociations remains to be established, but they are consistent with our theoretical analyses and with known physiological properties of cortical networks. Alpha- and beta-band activity have long been associated with inhibitory processing and are typically reduced during cortical activation (Klimesch et al., 2007; Pfurtscheller et al., 1996). Consistent with our simulations, these observations suggest that inhibitory interactions contribute to the divergence between phase propagation and information flow observed in vivo. A second notable finding was the consistency of DIFF across frequency bands. Whereas phase-based traveling waves exhibited marked frequency-dependent differences, DIFF revealed highly similar patterns across alpha, beta and broadband gamma in both marmosets and humans. In marmosets, these patterns consistently identified a source near the posterior parietal/dorsal visual cortex, with directed interactions predominantly toward posterior cortical regions, suggestive of feedback communication. The localization of alpha- and beta-band sources is broadly consistent with previous reports (Hoffman et al., 2024; Vezoli et al., 2021; Zhigalov and Jensen, 2020), whereas the broadband gamma pattern differs from prevailing accounts (Bastos et al., 2020, 2015). These findings suggest that frequency bands exhibiting distinct phase organizations may nevertheless participate in similar large-scale patterns of directed communication. More broadly, DIFF offers a complementary perspective on the relationship between oscillatory frequency and cortical hierarchy. In human EEG, theta —not alpha— displayed the strongest directed influences from frontal toward posterior regions, consistent with proposals that theta rhythms mediate long-range feedback interactions and are prominent in higher-order cortical networks (Cavanagh and Cohen, 2022; Cavanagh and Frank, 2014; Solomon et al., 2019; Vinck et al., 2023; Voloh et al., 2015; Womelsdorf et al., 2010a,b). By contrast, alpha activity was characterized by a prominent central source with widespread outward influences, consistent with proposed parietal generators of alpha oscillations (Bollimunta et al., 2008; Halgren et al., 2019; Zhigalov and Jensen, 2020) and potentially with activity in the default mode network (Menon, 2023; Raichle, 2015). Together, these observations suggest that directed information flow and oscillatory phase provide complementary views of large-scale cortical organization, and that integrating both may refine current models of feedforward and feedback communication.

### Complementary of Granger Causality and phase-based analyses

Granger Causality analysis has been widely used in the neurosciences for many applications (Brovelli et al., 2015, 2004b; Cekic et al., 2018; Ding et al., 2006; Marinazzo et al., 2011, 2008; Seth et al., 2015; Shojaie and Fox, 2022; Stokes and Purdon, 2017); however, to the best of our knowledge, Granger-causality has not been applied to analyze complex spatiotemporal patterns. When applied to a linear system, Granger causality provides a well-defined solution with uniquely separable contributions (Marinazzo et al., 2008; Seth et al., 2015) and reliably captures the causal directionality of input propagation into the system. We argue that Granger causality can capture the ground truth of causal influences for a linear system of coupled stochastic processes, in which causal influences across nodes can be uniquely decomposed, and the concept of causal influence has a well-established mathematical definition (Granger, 1969). Evidently, the phase pattern provides complementary information as an emergent phenomenon that is functionally relevant, but the two approaches may yield distinct yet complementary insights, as shown in the analysis of different datasets (Figure 5). For example, whereas the DIFF method provides directional information flow, averaged over several trials, the phase pattern, that can be extracted in single trials (Muller et al., 2014; Rubino et al., 2006), might encode information, sculpt the interactions or plasticity between nodes, and predict behavior (Benigno et al., 2023; Davis et al., 2020). The phase can also provide information about the speed of a wave traveling in a given direction. It is also important to emphasize that Granger-based methods have several limitations in general. First, uncorrelated or correlated noise can strongly affect Granger-causality inferences. For this problem, several solutions have been proposed, e.g., time-reversal controls (Haufe et al., 2014; Vinck et al., 2015) or Kalman-filtering approaches (Nalatore et al., 2007). Second, Granger causality is explicitly defined for stochastic time series and cannot therefore be used to infer causality in deterministic systems. Topological methods may, in such cases, allow for a definition and measurement of causality (Sugihara et al., 2012). Third, if a system is strongly nonlinear, nonlinear methods such as Transfer Entropy are required to capture causal relationships (Schreiber, 2000). In this case, the same framework and methodology can be applied using Transfer Entropy. Alternatively, more direct modeling approaches to capture the underlying dynamical system, e.g., dynamic causal modeling, could be suited for such cases. More generally, macroscopic traveling waves can emerge from an intricate propagation of activity in a complex underlying network (Budzinski et al., 2023), whose mechanistic origin may be hard to infer. That is, the mechanistic origin of phase gradients at the macroscopic level may be much more complex than the simple propagation of a single perturbation, e.g., axonal propagation. In addition, given the widespread connectivity in cortex (Vezoli et al., 2020), identifying causal relationships may require a multivariate approach. Fourth, Granger causality is not designed to be applied to a single trial but rather to a set of multiple trials, as it relies on fitting an underlying generative model. By contrast, phase-based methods can be applied to unique trials. It is possible, however, that distinct patterns of traveling waves can be detected across different trial sets and subsequently analyzed separately using Granger causality analyses (as shown in Figure 5). In this way, phase-based and Granger-causality measures can be applied synergistically.

Discrepancies between Granger causality and phase-based analyses may also arise from the unique properties of electrophysiological data. Importantly, the sign of LFP/EEG electrophysiological data is arbitrary and depends on the reference, and is subject to dipole reversals (Pesaran et al., 2018). Hence, the phase of electrophysiological signals depends on referencing schemes and the underlying generative model of the electric field. Granger-causality, on the other hand, is invariant to a scaling of signals and sign reversal. Thus, the DIFF analysis can, in principle, also be applied to current-source-density data or bipolarly derived data, corroborating the complementarity of the two approaches.

### Conclusion and Perspectives

Our results integrate theoretical, computational, and experimental approaches to demonstrate that the direction of traveling-wave propagation and the direction of information flow need not coincide. Although these two descriptions converge under specific conditions, they can diverge in cortical networks shaped by bidirectional connectivity, heterogeneous transmission delays, and excitatory–inhibitory interactions (Dugué and Chavane, 2025). By transforming Granger causal influences into a spatial vector field, DIFF provides a complementary framework for characterizing traveling waves in terms of directed neural communication while preserving the rich spatiotemporal information captured by phase-based analyses. These findings suggest that cortical traveling waves emerge from the interplay of multiple dynamical processes rather than from a simple propagation of activity across neural tissue. Integrating phase-based and causal descriptions may therefore provide a more complete understanding of largescale cortical communication and its role in neural computation. Extending this framework to nonlinear measures of directed interactions and more realistic models of cortical dynamics represents an important next step toward a unified description of information flow and wave propagation in the brain.

## Methods

### General VAR models

Consider a *k*-variate VAR model of order *T*, described by the system of discrete-time equations:

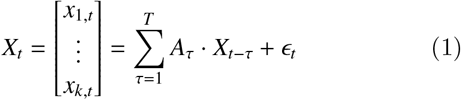

Here, *X*_*t*_ is a vector describing the evolution of a set of k variables. *A*_*τ*_ represents a (*k* × *k*)-coefficient matrix that describes the influence of the past state at lag *τ* (*X*_*t* − *τ*_) on the current state. The innovation term *ϵ*_*t*_ is a vector of the same dimension as *X*_*t*_ where each component is assumed to be white noise with zero mean and covariance matrix Σ.

Next, consider a simplification of this system, in which the nodes are sequentially organized, such that there are only direct interactions from a node *i* with the nodes *i* − 1 and *i* + 1. We call the flow from node *i* to node *i* + 1 “Direction 1” (D1), which is the information flow caused by the original perturbation. For a biological system, this may be either the forward connection (e.g., a sensory stimulus) or a feedback connection (e.g., imagery). We call the flow from node *i* + 1 to node *i* “Direction 2” (D2). In the system analyzed, ∀ *τ* ∈ [1, *T*], *A*_*τ*_ is a tridiagonal matrix with a specific set of coefficients defined as:

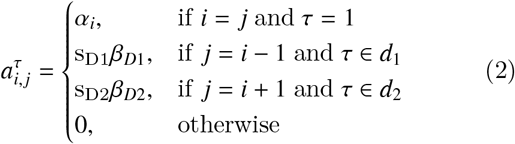

where α_*i*_, β_*D*1_, β_*D*2_ ∈ (ℝ^+^)^3^ are random variables drawn from N_trunc+_(0, 2), a Gaussian distribution truncated for all values below zero, s_D1_, s_D2_ ∈ {− 1, 1}^2^ determine the sign of the two opposite directions. and *d*_1_ and *d*_2_ are sets of delays that define connectivity at specific lags. Note that there is a unique weight, β_1_, connecting neighboring nodes in direction D1. Similar connectivity applies for D2, but the values of α_*i*_ depend on the nodes.

If we take a 2-variate VAR model, this yields in total 4 possible cases illustrated in Figure 1A, conditional on the value of s_D1_ and s_D2_. In the following, we will first consider the simplest VAR(1) case. Then, we will introduce models with higher orders, i.e., from VAR(2) to VAR(4), and with asymmetric delays. After a section describing a biologically plausible cortical model, we extend our results from 1D to 2D cases.

### The VAR(1) case

Starting from the general case described above, we first consider the condition with *d*_1_ = *d*_2_ = 1, i.e., using only the previous state to describe the current one :

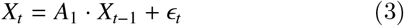

In this case, we test models with k=3 or k=7 nodes. For each of the four cases described above (Figure 1A), we run 1000 simulations of 500 time steps each. In each simulation, we determine the stability of the model by assessing that all eigenvalue modules associated with the companion matrix were |λ_*i*_ | < 1. We discard all runs in which the system was unstable.

The first node is fed an input signal, a series of independent, pseudo-random values drawn from a Gaussian distribution *N* (0, 1). In all nodes, we introduced independent Gaussian noise *ϵ*(*t*) from *N* (0, 0.1).

For all analyses, we only considered stable models. We first assessed the stability of the 1000 randomly generated models for each of the four conditions. We found that around 35% of simulations were stable for *k* = 3 and 25% for *k* = 7 for the four conditions.

### From VAR(1) to VAR(4)

We also consider the case with multiple temporal delays in a VAR model with order *p* > 1 (as in Equation 1).

Specifically, we investigated different conditions with specific sets of symmetric delays (i.e., *d* = *d*_1_ = *d*_2_) for information transfer between nodes. We study 3 different VAR models with *d* = {1, *δ*}, where *δ* ∈ {2, 3, 4}. In addition, we also considered the case where *d* = {1, 2, 3, 4}. The order of these VAR processes is given by max(*d*)

All the other parameters (i.e., number of simulations, time-steps, nodes, noise, and stability assessment) were the same as in the case without temporal delays. For all simulations, we assessed the stability of the 1000 simulations we performed with randomized coefficients. Overall, around 20% and 15% of the simulations were stable for *k* = 3 and *k* = 7, respectively. When considering several delays at once (*d* = {1, 2, 3, 4}), the stability dropped to less than 10% of the simulations, leaving, however, more than 50 runs in the lowest cases (cases 2 and 4). As previously, we considered only stable models for the analyses.

### Asymmetric Delays

We also study models with asymmetric delays (i.e., *d*_1_ ={*δ*_1_}, *d*_2_ = {*δ*_2_} and *δ*_1_ =f *δ*_2_). We simulate all combinations of models with delays varying from 1 to 100. Because of long delays, we want to run the phase-based and the Granger analysis once all nodes have been perturbed by the input, i.e., after time *t* = (*d*_1_ + *d*_2_) ∗ *k, k* being the number of nodes. For this reason, we increased the simulation length from 500 to 2000 time steps- and keep only the relevant part of the simulation. The results regarding both the phase-based and the Granger-based analysis were computed by taking the median of the phase/Granger values differences across all nodes for 50 stable simulations in each case.

### Simulations for a predictive coding model

We also apply our analysis to a biologically plausible model. Based on the generic VAR model given in Equation 1, we can introduce a specific set of parameters to match a given connectivity matrix or other existing models. Here, we set the parameters to match the dynamical system described in Alamia and VanRullen (2019).

We started from the equations of a two-layer system as described in (Alamia and VanRullen, 2019):

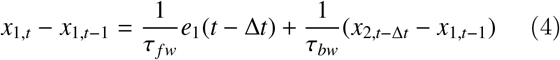

where *τ* _*fw*_ and *τ*_*bw*_ are time constants, *e*_1_(*t*) is the prediction error in the first layer:

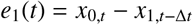

and *x*_0,*t*_ is the bottom-up input to the network at time *t*. This, eventually, simplifies to:

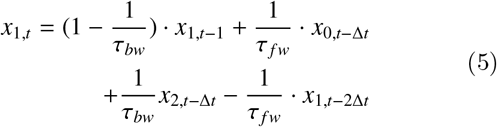

For this dynamical system, we have an equivalent VAR(2Δ*t*) model where all coefficient matrices have zero values except for *A*^1^, *A*^Δ*t*^, and *A*^2Δ*t*^. Thus, we have:

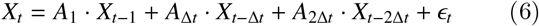

To match the dynamical system given by Equation 4, we need *A*_1_ to be a diagonal matrix with the diagonal coefficients equal to 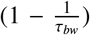, *A*_2Δ*t*_ to be another diagonal matrix with the diagonal coefficients equal to 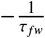 and *A*_Δ*t*_ to be a tridiagonal matrix with 0 on the diagonal,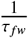 on the lower diagonal and 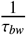 on the upper diagonal. As in Alamia and VanRullen (2019), we set *τ* _*fw*_ = 30 ms and *τ*_*bw*_ = 200 ms, these values granted a stable behavior and can be interpreted from a biological perspective as time constants. From this VAR(2Δ*t*) model, one can compute the frequency and oscillatory behavior, which depend on its parameters *τ* _*fw*_, *τ*_*bw*_ and Δ*t*, as demonstrated in Alamia and VanRullen (2019). Note that Equation 6 describes the 2-layer model given by Equation 5 but does not hold any assumption on the dimensionality of X. It can be easily generalized to *k* nodes by increasing the size of vector *X* and of the coefficient matrices *A* with the same structure as the one described in this case.

Considering this implementation, we tested models having *k* = 4 nodes. For each of the four cases described above (Figure 1A), we run 300 simulations of 500-time steps with independent Gaussian noise *ϵ*(*t*).

### Quantifying waves’ direction of propagation

In all simulations in which the model was stable (i.e., all eigenvalues |λ_*i*_ | < 1), we used two different approaches to quantify the activity’s direction of propagation. The first method was based on the estimates of the signal’s instantaneous phase; the second one applied Granger Causality to assess the flow’s direction.

### Phase-based analysis

At the end of each simulation, we computed the Hilbert transform *H*(*x*_*k*_) of each signal *x*_*k*_ for each *k* node and calculated the complex vector obtained by computing the phase difference between subsequent nodes at any time point *t*:

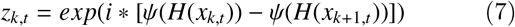

where ψ(*H*(*x*_*k,t*_)) is the phase of the Hilbert-transformed signal *x*_*t*_ at the node *k*. We then sum all complex vectors over time and compute the average phase difference between nodes:

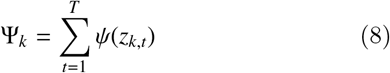

The sign of Ψ_*k*_ provides the phase shift between subsequent nodes: if positive, it indicates a wave propagating from node *k* to node *k* + 1; vice versa, a negative value reveals activity propagating from the higher to the lower nodes.

### Granger analysis

We used the multivariate Granger causality analysis (MVGC) toolbox in MATLAB (MathWorks) to compute the Granger causality between subsequent nodes in all cases (Barnett and Seth, 2014). For each condition and stable model, we first fit the model’s parameters, considering each simulation as a trial. We estimated the best model’s order by optimizing the Akaike Information Criterion (we obtained similar results using the Bayesian Information Criterion). For the 1-D VAR(1) case, we fitted the model’s order for k=3 and k=7 nodes, optimizing Akaike’s criterion, and we obtained a fit of the third and fifth order, respectively, for all conditions. The results reported did not change qualitatively when the model’s order was set to equal 1 before fitting the model’s parameters.

Besides estimating the model’s parameters, such a fit provides the covariance matrix of the residuals, which is next used to calculate the autocovariance sequence according to the estimated VAR model. Lastly, the pair-wise time-domain Granger causalities are computed for each pair of *x*_*i*_ from the covariance sequence. Significant effects were estimated against the null hypothesis of zero causality based on a theoretical asymptotic null distribution (Barnett and Seth, 2014).

### Generalization to 2D networks

#### Nodes configuration and simulations

We generalized our results from the one-dimensional case, i.e., a series of consecutively connected nodes, to the two-dimensional case, in which the nodes are organized in a square grid and connected to their neighbors. We considered squared grids of side *k* × *k*, with *k* = 9. It is possible to formalize the 2D grid arrangement as a 1D case with long-range connections (i.e., between non-subsequent nodes). As specified in Figure 1C, the parameters connecting node *i* to node *i* − *k* determine D1, whereas parameters from node *i* to node *i* + *k* determine D2. The parameters between node *i* and its neighbors *i* ∓ 1 characterize the lateral connections. As in the 1D case, we can fully describe the model via the parameters of the matrix *A*_*τ*_. For this 2D case, we focus on a VAR(1) process, and we can describe *A*_1_ as:

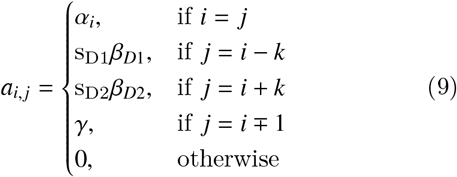

where α, β_1_, β_2_, γ ∈ (ℝ^+^)^4^ are random variables drawn from *N*_trunc+_(0, 2), a Gaussian distribution truncated for all values below zero, and s_D1_, s_D2_ ∈ {−1, 1}^2^ determine the sign of the two opposite directions. Considering this configuration, we tested 300 simulations of 100 time-steps in which we perturbed a subset of specific nodes either at different locations, or at the lower edge of the grid, with a series of independent, pseudo-random values drawn from a Gaussian distribution *N*(0, 2). In all cases, we introduced independent Gaussian noise E(*t*) from *N*(0, η) at each node. The value of η was modulated to modulate the noise levels over different simulations.

#### Phase-based analysis

We performed the phase-based analysis by estimating a phase-based vector field in each grid. In particular, for all nodes *k* we computed the phase differences between the Hilbert transforms of each signal *H*(*x*_*k*_) and the neighboring nodes whose distance to node *k* was smaller than or equal to 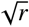. To consider the circle capturing direct diagonal neighbors as well as directly connected neighbors, we set *r* = 2. Such a phase difference was defined as:

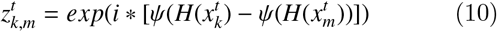

where 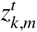 is the phase difference between the *k* node and its neighbor *m* at time *t*. For all neighbors *m*, we computed the sinusoid of such phase difference averaged across time points:

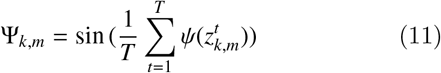

Lastly, we average the phase difference Ψ_*k,m*_ over all neighbors *m*, considering their relative position with respect to the node *k*. This results in a vector field in which each vector, centered at every node, points to the mean phase difference between its neighbors whose distance to *k* is smaller than or equal to 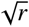.

#### DIFF method

Similar to the phase-based vector field, we computed a vector field based on the directional information flow estimated via Granger measures. The Directional Information

Flow Field (DIFF) is then a vector field obtained by computing the pair-wise Granger causality between each node and its neighbors whose distance to *k* was smaller than or equal to 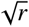 (as before, we set *r* = 2). The contribution of each neighbor *k* to a given node *i* is computed as the difference between the Granger causality from node *i* to *k* and the Granger causality from *k* to *i*.

For every node *k*, we averaged the Granger values over all neighbors *m*, considering their relative position with respect to the node *k*. This generates a vector field *GC*_*x,y*_ where, at each node’s position *x, y*, the vector points in the averaged information flow estimated as the mean Granger value between its neighbors. As above, all Granger analyses were computed using the MVGC toolbox in Matlab (Barnett and Seth, 2014). Finally, for both the phase-based and the Granger-based vector fields *VF*_*x,y*_, we estimated the absolute divergence at every node as:

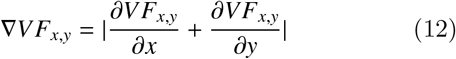

Crucially, values of *VF*_*x,y*_ larger than 0 locally identify the source of the external perturbations.

#### Electrophysiological data

Two previously published datasets were used: ECoG recordings marmosets (Canales-Johnson et al., 2021; Gelens et al., 2024), and one scalp-EEG dataset in healthy human participants (Pang et al., 2020). For all datasets, phase-based and Granger vector fields were computed in the same manner as for the 2D signal simulations. Importantly, we determine the size of the radius to consider neighboring electrodes to have around 10% of the electrodes, corresponding on average to 5.3 neighboring electrodes for the marmosets, and 7.2 for the scalp EEG (see Figure S4A). Below, we briefly describe the minimal preprocessing steps for each dataset. We refer to the original publications for further details.

#### ECoG data on marmosets

These data were recorded in two marmosets (i.e., GO [64 electrodes], and FR [32 electrodes]) performing a roving oddball paradigm (Canales-Johnson et al., 2021; Gelens et al., 2024), in which trains of identical tones (3, 5, or 11 repetitions) were presented continuously. The first tone of each train served as an unexpected deviant, and the last tone as an expected standard. Pure tones of varying frequencies were presented with fixed durations and interstimulus intervals, and only the final standards were analyzed to balance standard and deviant counts. ECoG signals were recorded at 1 kHz using a multichannel system (0.3–500 Hz bandpass), rereferenced to the average, high-pass filtered at 0.5 Hz, and segmented into standard and deviant epochs (−400 to 400 ms relative to tone onset; 1440 standard and 1440 deviant epochs for GO, and 720 standard and 720 deviant epochs for FR).

#### Scalp EEG on humans

The EEG data were recorded while 13 healthy subjects performed a visual detection task. Participants completed five sessions, each comprising 10 experimental blocks of six trials. Each trial consisted of 5 seconds of visual stimulation (stimulus ON condition) followed by 5 seconds of blank screen (stimulus OFF condition), while subjects maintained fixation and covertly attended the stimulus to detect a brief luminance-decrement target. Continuous EEG was recorded using a 64-channel active BioSemi system at 1024 Hz, with three additional ocular electrodes. Preprocessing was performed in EEGlab using custom scripts, including both target-present and target-absent trials. Noisy channels were rejected; the data were downsampled to 160 Hz; line noise was removed using a 47–53 Hz notch filter; signals were rereferenced to the average; and slow drifts were removed with a high-pass filter with a cutoff frequency greater than 1 Hz. Data containing eye movements, blinks, muscle artifacts, and contaminated epochs were rejected.

## Code availability

The code to replicate the analysis is available at https://github.com/artipago/comparing-phase-based-and-Granger-based-analyses.

## Author contribution

AA, FC, and MV conceived of the study. AA and AG analyzed data. AA and MV wrote the paper, with comments from the other authors.

## Acknowledgements

The authors wish to thank Lyle Muller and Laura Dugué for their helpful and constructive comments. MV was supported by an ERC starting grant (850861) SPATEMP; DFG VI Grants (908/5-1 and 908/7-1); an NWO VIDI Grant; and the Dutch Brain Interface Initiative (DBI2). AA was supported by ERC starting grant (OSCI-PRED, no. 101075930). ACJ is funded by an ANID/FONDECYT Regular (1240899) and ANID/FONDECYT Regular (1251273) research grants, and by a Swedish Research Council Project grant (VR; 2025-03245). Views and opinions expressed are, however, those of the authors only and do not necessarily reflect those of the European Union or the European Research Council (ERC). Neither the European Union nor the granting authority can be held responsible for them.

## Supplementary Materials

### Supplementary: All simulations with 1D networks

Below, we describe the results for all the simulations in the one-dimensional case. We generalized and replicated our results to VAR(k) models, where *k* ∈ {2, 3, 4}, and to a cortical model grounded in the hierarchical predictive model of inter-areal interactions (Alamia and VanRullen, 2019), which includes biologically plausible temporal delays between subsequent cortical nodes.

### From VAR(1) to VAR(4)

We tested whether the effect could be generalized to cases with different sets of temporal delays. First, we assessed the stability of the 1000 simulations we performed with randomized coefficients. Overall, around 20% and 15% of the simulations were stable for *k* = 3 and *k* = 7, respectively. When considering several delays at once (*d* = {1, 2, 3, 4}), the stability dropped to less than 10% of the simulations, leaving, however, more than 50 runs in the lowest cases (cases 2 and 4). As previously, we considered only stable models for the subsequent analyses.

### Phase-based analysis

As in the case without temporal delays, we found consistent results for k=3 and k=7 nodes (see Figure ??A,B for the VAR(2) case). In particular, we found a backward propagation in the first and fourth cases, as confirmed by a negative phase difference between all subsequent nodes, but the opposite direction in the second and third cases, corroborating our previous results. These results were very consistent across the different delays. As in the VAR(1) case, a Von Mises test (V-test) for non-uniformity of circular data considering the mean direction in any pair of nodes confirmed the results: in all conditions and pair of nodes, we obtained for k=3 all V-value > 120 and *p* < 0.0001 when considering *τ* = [1, 4], and V-value > 65 and *p* < 0.0001 when having several delays. Similarly, for k=7 and *τ* = [1, 4], we obtained all V-value > 50 (with larger values ≈ 110 for lower nodes, but invariably significant) and *p* < 0.0001, and V-value > 20 when considering several delays.

As in the case without temporal delays, the results of our simulations corroborated the finding that a positive sign from the node *i* to node *i* + 1 produces a phase difference consistent with D1 direction (i.e., a forward flow from the input), whereas a negative one leads to a phase difference corresponding to a flow in D2 direction (towards the input).

### Granger analysis

The Granger analysis was applied to all nodes’ activity to estimate the causal propagation direction. We fitted the model’s order for k=3 and k=7 nodes for each temporal delay, optimizing Akaike’s criterion. For a single temporal delay, i.e., *τ* ∈ {2, 3, 4}, we obtained orders between 3 and 5, and we obtained an order equal to 8 when considering all the temporal delays, i.e., *τ* = {1, .., 4}.

Considering the k=3 case, we found very similar time-domain Granger Causality (GC) values when having a single temporal delay. For instance, when *τ* = 2, we found 1.377 ± 0.10 from node 1 to node 2, and 0.88 ± 0.20 from node 2 to node 3, and 0.284 ± 0.04 from node 1 to node 3. All other GC values were smaller than 0.05. When considering the case with all temporal delays, we obtained relatively smaller values: 0.804 ± 0.04 from node 1 to node 2, and 0.56 ± 0.05 from node 2 to node 3, and 0.10 ± 0.02 from node 1 to node 3.

Regarding k=7 and *τ* = 2 (very similar values were observed in the other cases having one single temporal delay), we found the larger GC values in lower nodes: progressively from node 1 to node 7 we reported 1.433 ± 0.31, 0.768 ± 0.11, 0.16 ± 0.06, and then all other values were smaller than 0.07 ± 0.03. Similarly, when *τ* ={1, .., 4}, we found value decreasing from lower to higher nodes: 0.876 ± 0.08, 0.346 ± 0.03, 0.097 ± 0.2, and then all other values were smaller than 0.05 ± 0.02.

Altogether, the Granger analysis confirms that the causal flow of information propagates the input information from the perturbed node 1 to other nodes (Direction D1), irrespective of the sign of the *a*_*i,i*+1_ coefficient.

### Simulations for a predictive coding model

Next, we generalized our results to a model grounded in the predictive coding framework (Alamia and VanRullen, 2019), which includes biologically plausible temporal delays between subsequent nodes. We simulated 300 instances of the model for 500 time steps for each of the four conditions. The model had *k* = 4 nodes, and its parameters were chosen to be biologically interpretable and with a stable behavior (see Methods, and (Alamia and VanRullen, 2019; Schwenk and Alamia, 2024)).

### Phase-based analysis

The results based on the phase analysis support the same conclusions as in the previous simulations, as shown in Figure ??C. In particular, we observed opposite propagation directions depending on the sign of the *a*_*i,i*+1_ coefficient. In the first and fourth cases, when the *a*_*i,i*+1_ coefficient is negative, we observe a consistent, negative phase difference across subsequent nodes, revealing a propagation along direction D2. On the other hand, a positive *a*_*i,i*+1_ coefficient (i.e., second and third cases) generates the opposite direction D1, as confirmed by a positive phase difference between subsequent nodes. These results were confirmed by a V-test for non-uniformity of circular data performed over the 300 simulations and considering the mean direction in any pair of nodes (in all conditions and pair of nodes V-value> 280 and *p* < 0.0001).

### Granger analysis

We then estimated the Granger causalities between all subsequent nodes. We assessed the best model’s order for each condition, having an order of 19 for the first condition and 18 for all the other ones. Figure ??C illustrates the time-domain Granger Causality (GC) values between all nodes. Asterisks reveal a significant effect when compared against a theoretical asymptotic null distribution (Barnett and Seth (2014)). Across conditions, we found decreasing GC values across the hierarchy: 1.407 ± 0.36 from node 1 to node 2, 0.340 ± 0.03 from node 2 to node 3, and 0.05 ± 0.01 from node 3 to node 4. All other GC values (i.e., in the opposite direction and between non-subsequent nodes) were smaller than 0.02 ± 0.01.

### Supplementary Figure 01: All simulations with 1D networks

#### Supplementary simulations with Planar 2D traveling waves

As reported in the main text, we also investigate the case of a planar wave propagating from the lower edge of the network (akin to node 1 in the 1D case), as shown in Figure S3. Similarly to the case with localized sources generating radial waves, a planar wave can also be reliably identified by the DIFF method applied to the Granger-based vector field, irrespective of the amount of noise level E (S3C). Different from the Granger-based vector field, the divergence of the phase vector field does not identify the source of the planar wave in all cases.

**Figure S1:**
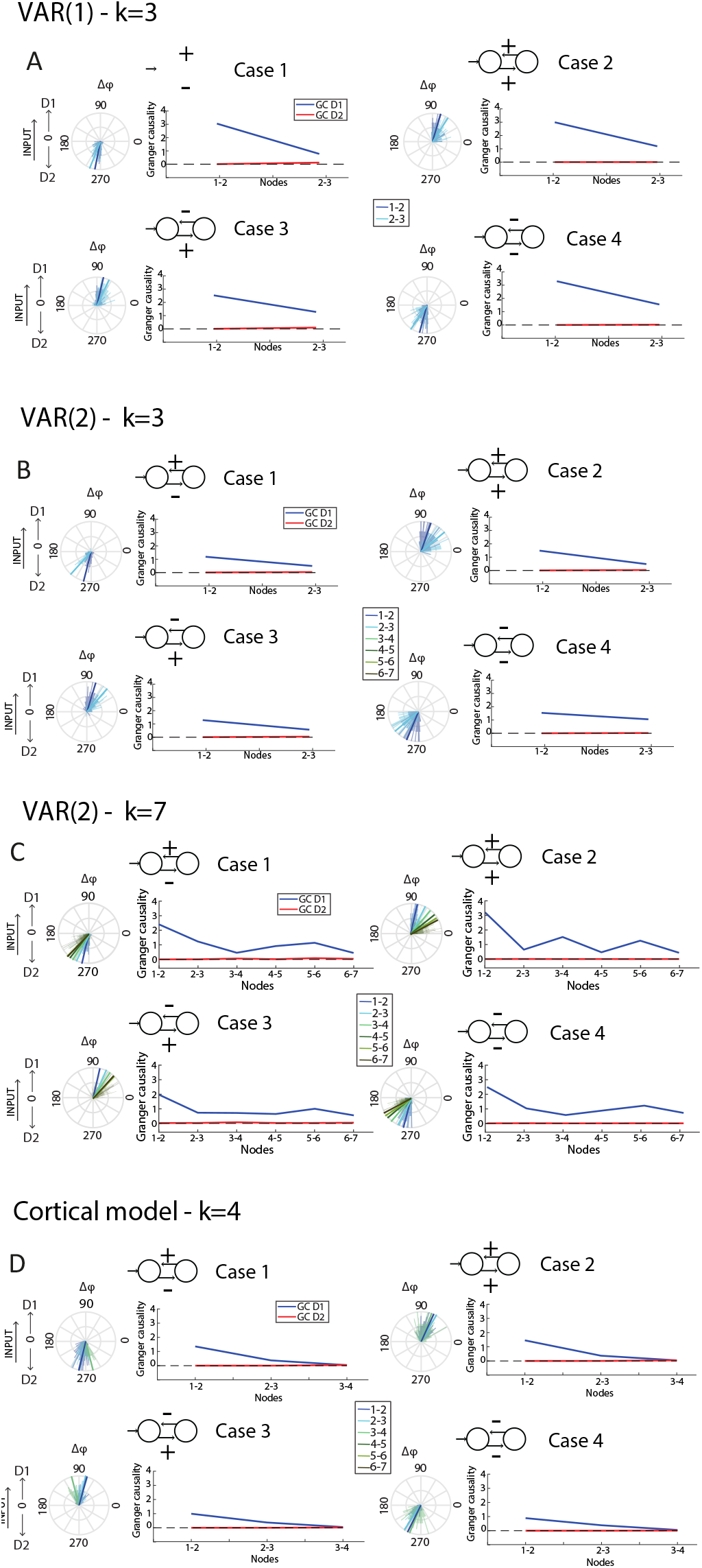
Results for the VAR(2) condition with k=3 nodes (panel A) and k=7 nodes (panel B). Panel C shows the results for the cortical relevant model (based on predictive coding dynamics). Each panel shows the four cases, as specified in the upper insert. For each case, a rose plot illustrates the average phase difference in degrees (90° reveals a flow toward D1, 270° towards D2), and the plot on the right indicates pairwise Granger causality between subsequent nodes. In all panels and simulations, the results show that Granger values consistently reveal the expected directionality (i.e., D1), whereas phase differences consistently reveal opposite results in cases 1 and 4.

**Figure S2:**
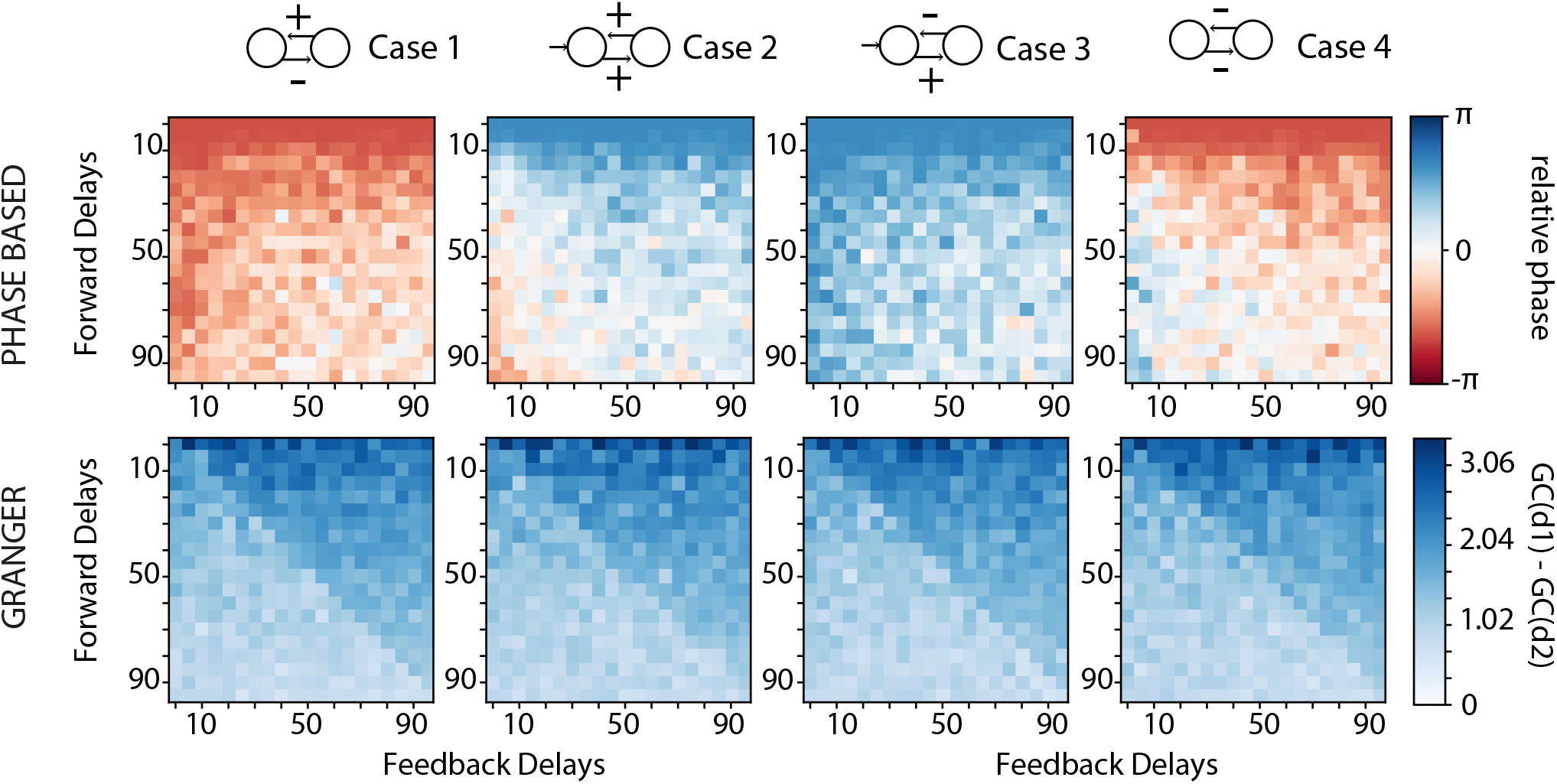
Results of the phase-based and Granger analyses for asymmetric delays. Each pixel corresponds to a pair of delays (*δ*_1_, *δ*_2_) for directions D1 and D2. The top row shows the median phase difference across nodes. For visualization purposes, we apply a tanh transformation for the visualization of the relative phases. Case 2 and Case 4 reveal a change of the phase sign when delays of D1 are larger than delays of D2. The bottom row shows the difference between the average of Granger causality coefficients in the two distinct directions (GC(D1)-GC(D2))

**Figure S3:**
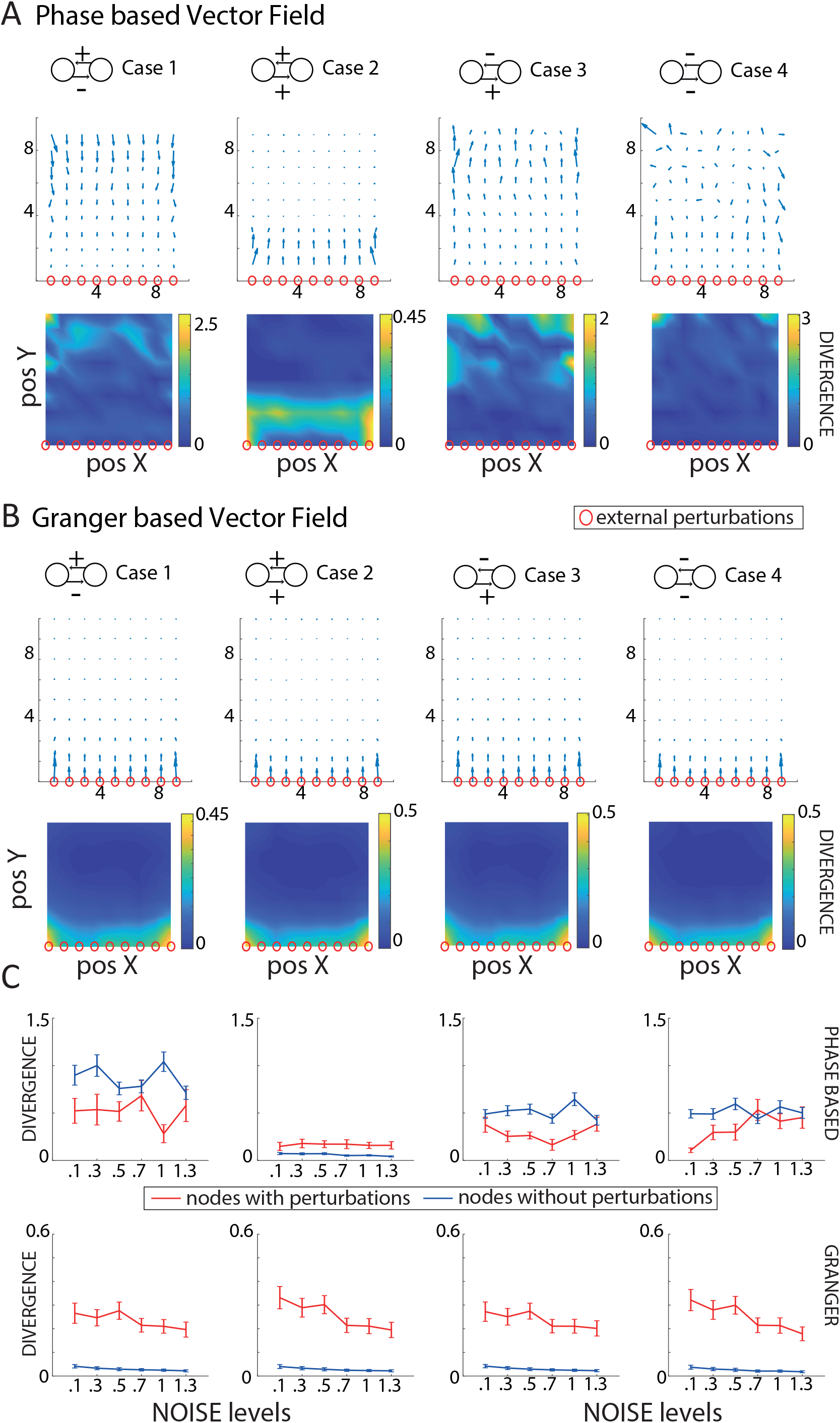
Results for the 2D networks when perturbing the lower edge of the network. Panels A show the results of the phase-based vector field for the four cases, as summarized above each column. The first line represents the vector field, and the second one its divergence. The red nodes are the sources of the external perturbations. Panel B shows the results for the Granger-based (i.e., DIFF) vector fields, same organization as in panel A. Panel C) shows the changes in the divergence as a function of the noise E introduced independently in all nodes. Blue lines are the nodes without perturbation (mean ± standard deviations), whereas the red lines represent the divergence in the perturbed nodes.

**Figure S4:**
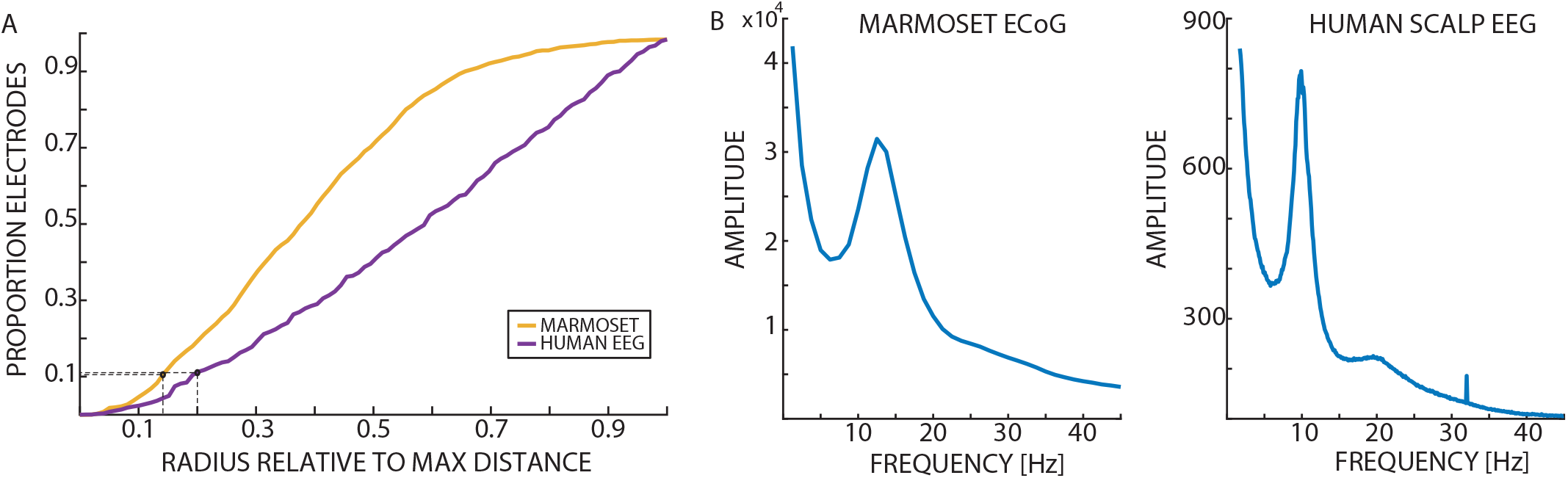
A) Proportion of neighboring electrodes as a function of radius. The figure shows the proportion of electrodes, relative to the total number of electrodes across all datasets, that were included in estimating the resultant vector for each electrode using both the DIFF and phase-based methods. The x-axis denotes the normalized radius, with the maximum inter-electrode distance used as the reference. Black circles indicate the radius selected for each dataset, corresponding, on average, to 5.3 neighbors for the marmoset and 7.2 for the scalp EEG. B) Power spectra for the Marmoset and EEG data, averaging over trials, channels, and (in the case of the EEG) participants.

**Figure S5:**
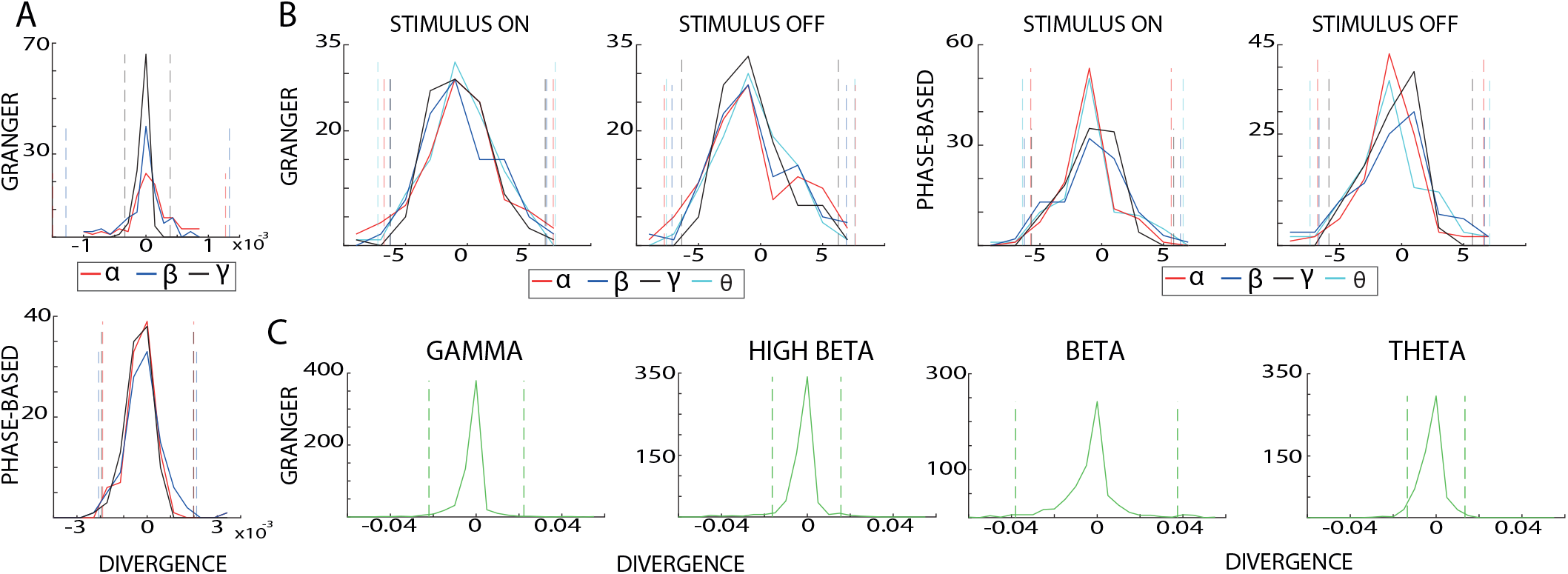
A) The plots show the distributions of divergence values for the marmoset across the three frequency bands (alpha in red, beta in blue, and gamma in black) for the Granger-based and phase-based analyses. B) Distributions of divergence values for the scalp-EEG recordings in humans, across the three frequency bands (theta in cyan, alpha in red, beta in blue, and gamma in black), for both stimulus ON and stimulus OFF conditions, for both methods. C) Divergence distributions for the macaque for each frequency band separately. In all plots, dashed lines show the 2.5 and 97.5 percentiles obtained on surrogate distributions for each frequency band.

**Figure S6:**
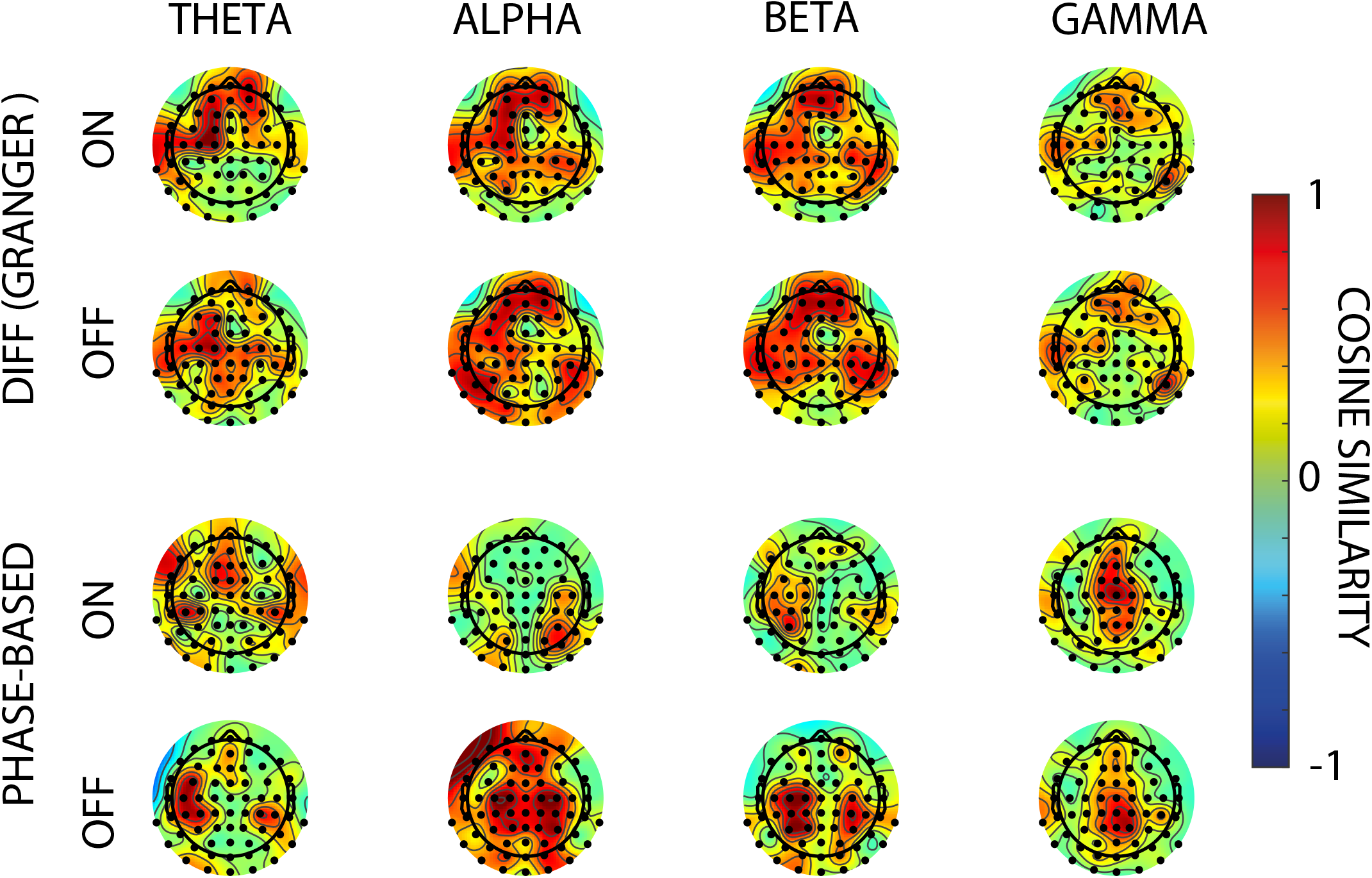
Scalp-EEG cosine similarity between participants. The figure illustrates the scalp topography of the average inter-participant cosine similarity of vector fields across all frequency bands for both methods. Using the DIFF method, we observe relatively high similarity between participants in all bands except the gamma band. The phase-based method also shows high inter-participant similarity during rest (i.e., the stimulus OFF condition), whereas similarity is reduced during visual stimulation (i.e., stimulus ON).

**Figure S7:**
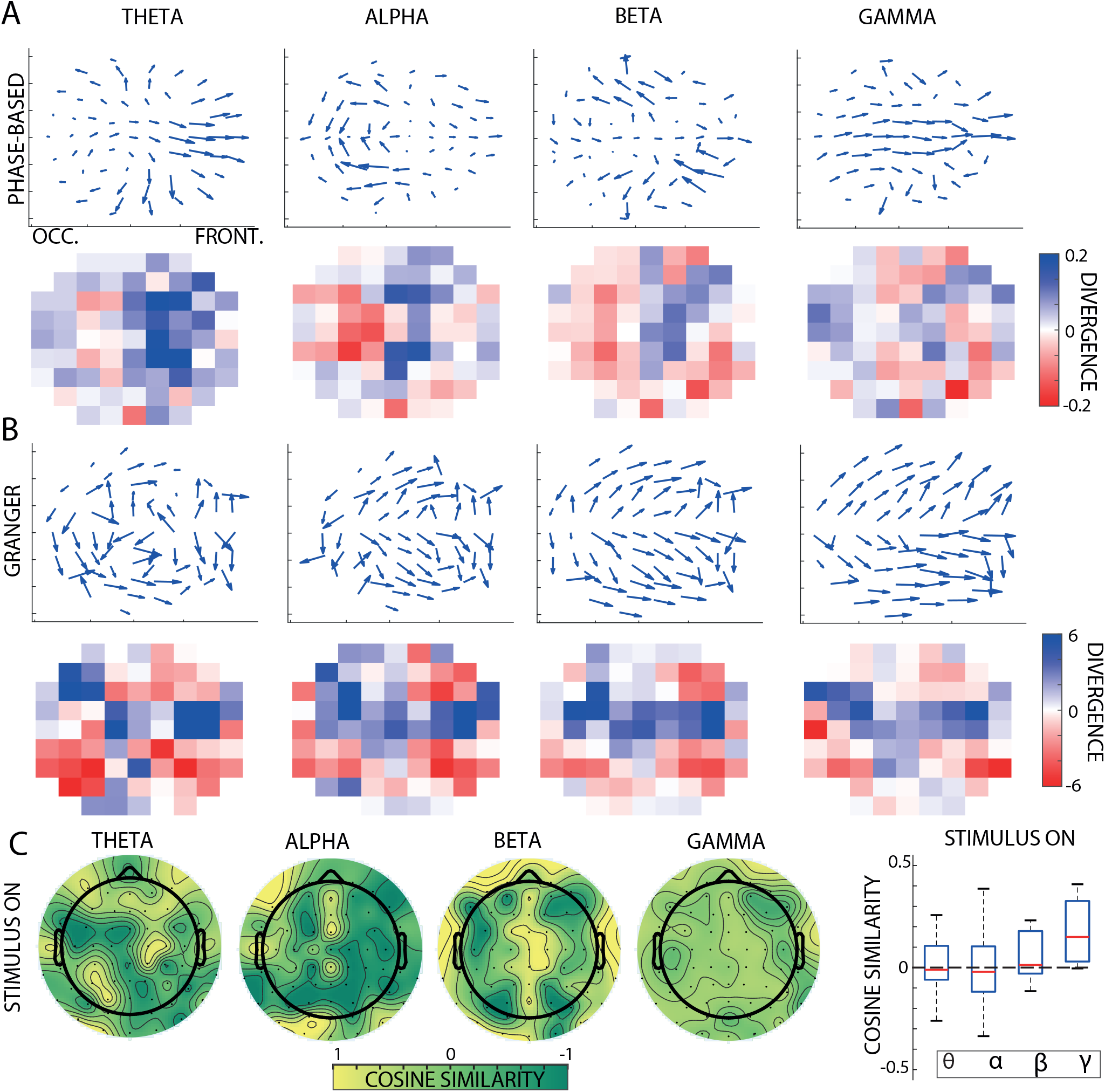
Results for the Scalp-EEG data during visual stimulation. Panels A and B show the average vector fields obtained using the DIFF and phase-based methods applied to scalp EEG data during visual stimulation (first row). The second row shows the average divergence of these vector fields across participants. Panel C presents the cosine similarity between the DIFF and phase-based vector fields. With the exception of the gamma band, the two methods exhibit relatively low similarity in occipital and parietal regions. The plot on the right shows the cosine similarity averaged across electrodes for each frequency band.

**Figure S8:**
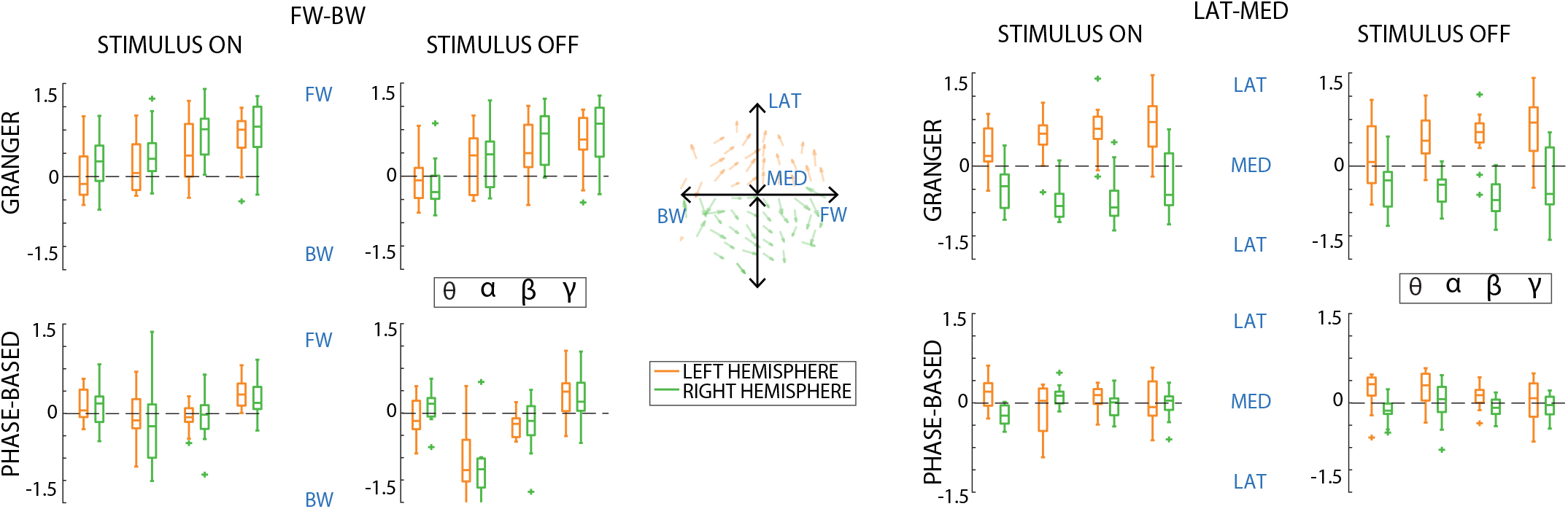
The figure shows the main direction of propagation for both methods, for all frequency bands, in scalpEEG recordings. We considered the average of the real and imaginary components of all vectors in each hemisphere separately (left in orange and right in green), representing the anterior-posterior and medio-lateral propagation, respectively. We observed significant differences between the two methods, especially in the medio-lateral axis.

**Figure S9:**
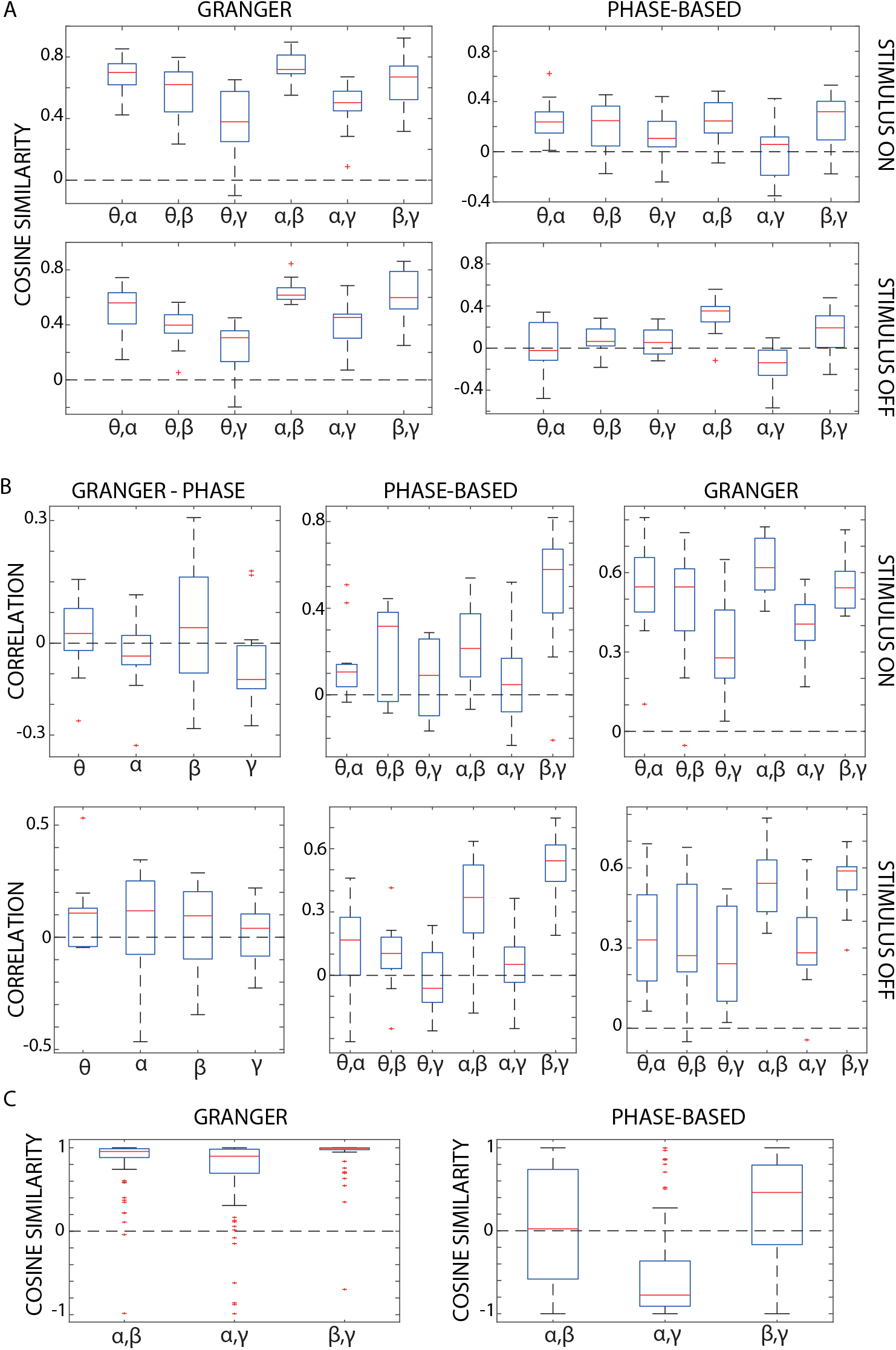
A) Each panel shows the boxplots of cosine similarity values across participants between vector fields estimated from different frequency bands using either the Granger-based method (left column) or the phase-based method (right column), during stimulus ON (top row) and stimulus OFF (bottom row) conditions. B) Boxplots of correlations between divergence maps across participants. The first panel compares divergence maps between Granger-based and phase-based vector fields. The remaining panels show correlations between frequency bands within each method (phase-based and Granger-based), separately for stimulus ON (top row) and stimulus OFF (bottom row) conditions. C) Cosine similarity between frequency-band-specific vector fields in the marmoset recordings, computed using the Granger-based method (left panel) and the phase-based method (right panel).

## References

Alamia, A., Terral, L., D’ambra, M.R., VanRullen, R., 2023. Distinct roles of forward and backward alpha-band waves in spatial visual attention. Elife 12, e85035.

Alamia, A., VanRullen, R., 2019. Alpha oscillations and traveling waves: Signatures of predictive coding? PLoS Biology 17, e3000487.

Alexander, D.M., Jurica, P., Trengove, C., Nikolaev, A.R., Gepshtein, S., Zvyagintsev, M., Mathiak, K., Schulze-Bonhage, A., Ruescher, J., Ball, T., et al., 2013. Traveling waves and trial averaging: the nature of single-trial and averaged brain responses in large-scale cortical signals. Neuroimage 73, 95–112.

Barnett, L., Seth, A.K., 2014. The mvgc multivariate granger causality toolbox: a new approach to granger-causal inference. Journal of neuroscience methods 223, 50–68.

Bastos, A.M., Lundqvist, M., Waite, A.S., Kopell, N., Miller, E.K., 2020. Layer and rhythm specificity for predictive routing. Proceedings of the National Academy of Sciences 117, 31459–31469.

Bastos, A.M., Vezoli, J., Bosman, C.A., Schoffelen, J.M., Oostenveld, R., Dowdall, J.R., De Weerd, P., Kennedy, H., Fries, P., 2015. Visual areas exert feedforward and feedback influences through distinct frequency channels. Neuron 85, 390–401.

Benigno, G.B., Budzinski, R.C., Davis, Z.W., Reynolds, J.H., Muller, L., 2023. Waves traveling over a map of visual space can ignite short-term predictions of sensory input. Nature Communications 14, 3409.

Benucci, A., Frazor, R.A., Carandini, M., 2007. Standing waves and traveling waves distinguish two circuits in visual cortex. Neuron 55, 103–117.

Bollimunta, A., Chen, Y., Schroeder, C.E., Ding, M., 2008. Neuronal mechanisms of cortical alpha oscillations in awake-behaving macaques. Journal of Neuroscience 28, 9976–9988.

Briggs, F., 2020. Role of feedback connections in central visual processing. Annual review of vision science 6, 313–334.

Brovelli, A., Chicharro, D., Badier, J.M., Wang, H., Jirsa, V., 2015. Characterization of cortical networks and corticocortical functional connectivity mediating arbitrary visuomotor mapping. Journal of Neuroscience 35, 12643–12658.

Brovelli, A., Ding, M., Ledberg, A., Chen, Y., Nakamura, R., Bressler, S.L., 2004a. Beta oscillations in a large-scale sensorimotor cortical network: directional influences revealed by granger causality. Proceedings of the National Academy of Sciences 101, 9849–9854.

Brovelli, A., Ding, M., Ledberg, A., Chen, Y., Nakamura, R., Bressler, S.L., 2004b. Beta oscillations in a large-scale sensorimotor cortical network: directional influences revealed by granger causality. Proceedings of the National Academy of Sciences 101, 9849–9854.

Budzinski, R.C., Nguyen, T.T., Benigno, G.B., Đoàn, J., Mináč, J., Sejnowski, T.J., Muller, L.E., 2023. Analytical prediction of specific spatiotemporal patterns in nonlinear oscillator networks with distance-dependent time delays. Physical Review Research 5, 013159.

Callaway, E.M., 2004. Feedforward, feedback and inhibitory connections in primate visual cortex. Neural Networks 17, 625–632.

Campbell, J.M., Davis, T.S., Anderson, D.N., Arain, A., Davis, Z., Inman, C.S., Smith, E.H., Rolston, J.D., 2024. Macroscale traveling waves evoked by single-pulse stimulation of the human brain. bioRxiv, 2023–03.

Canales-Johnson, A., Borges, A.F.T., Komatsu, M., Fujii, N., Fahrenfort, J.J., Miller, K.J., Noreika, V., 2021. Broadband dynamics rather than frequency-specific rhythms underlie prediction error in the primate auditory cortex. Journal of Neuroscience 41, 9374–9391.

Cavanagh, J.F., Cohen, M.X., 2022. Frontal midline theta as a model specimen of cortical theta, in: The Oxford handbook of EEG frequency. Oxford University Press. volume 178.

Cavanagh, J.F., Frank, M.J., 2014. Frontal theta as a mechanism for cognitive control. Trends in cognitive sciences 18, 414–421.

Cekic, S., Grandjean, D., Renaud, O., 2018. Time, frequency, and time-varying granger-causality measures in neuroscience. Statistics in medicine 37, 1910–1931.

Chemla, S., Roux, S., Reynaud, A., Chavane, F., VanRullen, R., 2019. Revealing α oscillatory activity using voltage-sensitive dye imaging in monkey v1. bioRxiv, 810325.

Cruddas, J., Pang, J.C., Fornito, A., 2026. Cortical traveling waves in time and space: Physics, physiology, and psychology. Neuron 114, 985–1005.

Das, A., Zabeh, E., Jacobs, J., 2023. How to detect and analyze traveling waves in human intracranial eeg oscillations?, in: Intracranial EEG: A Guide for Cognitive Neuroscientists. Springer, pp. 487–505.

Davis, Z.W., Muller, L., Martinez-Trujillo, J., Sejnowski, T., Reynolds, J.H., 2020. Spontaneous travelling cortical waves gate perception in behaving primates. Nature 587, 432–436.

Ding, M., Chen, Y., Bressler, S.L., 2006. Granger causality: basic theory and application to neuroscience. Handbook of time series analysis: recent theoretical developments and applications, 437–460.

Dowdall, J.R., Schneider, M., Vinck, M., 2023. Attentional modulation of inter-areal coherence explained by frequency shifts. NeuroImage 277, 120256.

Dowdall, J.R., Vinck, M., 2023. Coherence fails to reliably capture inter-areal interactions in bidirectional neural systems with transmission delays. NeuroImage 271, 119998.

Duan, P., Yang, F., Chen, T., Shah, S.L., 2013. Direct causality detection via the transfer entropy approach. IEEE transactions on control systems technology 21, 2052–2066.

Dugué, L., Chavane, F., 2025. Traveling waves across scales: Different mechanisms but same canonical computation? ELife 14, e106753.

Ermentrout, G.B., Kleinfeld, D., 2001. Traveling electrical waves in cortex: insights from phase dynamics and speculation on a computational role. Neuron 29, 33–44.

Freeman, W.J., Skarda, C.A., 1985. Spatial eeg patterns, non-linear dynamics and perception: the neo-sherringtonian view. Brain Research Reviews 10, 147–175.

Galas, L., Donovan, I., Dugué, L., 2025. Attention rhythmically shapes sensory tuning. Journal of Neuroscience.

Gelens, F., Äijälä, J., Roberts, L., Komatsu, M., Uran, C., Jensen, M.A., Miller, K.J., Ince, R.A.A., Garagnani, M., Vinck, M., Canales-Johnson, A., 2024. Distributed representations of prediction error signals across the cortical hierarchy are synergistic. Nature Communications 15. URL: http://dx.doi.org/10.1038/s41467-024-48329-7, doi:10.1038/s41467-024-48329-7.

Granger, C.W., 1969. Investigating causal relations by econometric models and cross-spectral methods. Econometrica: journal of the Econometric Society, 424–438.

Halgren, M., Ulbert, I., Bastuji, H., Fabó, D., Erőss, L., Rey, M., Devinsky, O., Doyle, W.K., Mak-McCully, R., Halgren, E., et al., 2019. The generation and propagation of the human alpha rhythm. Proceedings of the National Academy of Sciences 116, 23772–23782.

Hatsopoulos, N.G., Ojakangas, C.L., Paninski, L., Donoghue, J.P., 1998. Information about movement direction obtained from synchronous activity of motor cortical neurons. Proceedings of the National Academy of Sciences 95, 15706–15711.

Haufe, S., Meinecke, F., Görgen, K., Dähne, S., Haynes, J.D., Blankertz, B., Bießmann, F., 2014. On the interpretation of weight vectors of linear models in multivariate neuroimaging. Neuroimage 87, 96–110.

Hoffman, S.J., Dotson, N.M., Lima, V., Gray, C.M., 2024. The primate cortical lfp exhibits multiple spectral and temporal gradients and widespread task dependence during visual short-term memory. Journal of Neurophysiology 132, 206–225.

Isaacson, J.S., Scanziani, M., 2011. How inhibition shapes cortical activity. Neuron 72, 231–243.

Klimesch, W., Sauseng, P., Hanslmayr, S., 2007. Eeg alpha oscillations: the inhibition–timing hypothesis. Brain research reviews 53, 63–88.

Lefèvre, J., Baillet, S., 2009. Optical flow approaches to the identification of brain dynamics. Human brain mapping 30, 1887–1897.

Liu, X., Wiesman, A.I., Baillet, S., 2026. Hierarchical flows of human cortical activity. bioRxiv.

Lubenov, E.V., Siapas, A.G., 2009. Hippocampal theta oscillations are travelling waves. Nature 459, 534–539.

Luo, C., VanRullen, R., Alamia, A., 2021. Conscious perception and perceptual echoes: a binocular rivalry study. Neuroscience of Consciousness 2021, iab007.

Marinazzo, D., Liao, W., Chen, H., Stramaglia, S., 2011. Nonlinear connectivity by granger causality. Neuroimage 58, 330–338.

Marinazzo, D., Pellicoro, M., Stramaglia, S., 2008. Kernel-granger causality and the analysis of dynamical networks. Physical Review E—Statistical, Nonlinear, and Soft Matter Physics 77, 056215.

Menon, V., 2023. 20 years of the default mode network: A review and synthesis. Neuron 111, 2469–2487.

Mohan, U.R., Zhang, H., Ermentrout, B., Jacobs, J., 2024. The direction of theta and alpha travelling waves modulates human memory processing. Nature Human Behaviour 8, 1124–1135.

Muller, L., Chavane, F., Reynolds, J., Sejnowski, T.J., 2018. Cortical travelling waves: mechanisms and computational principles. Nature Reviews Neuroscience 19, 255–268.

Muller, L., Reynaud, A., Chavane, F., Destexhe, A., 2014. The stimulus-evoked population response in visual cortex of awake monkey is a propagating wave. Nature communications 5, 3675.

Murphy, P.C., Duckett, S.G., Sillito, A.M., 1999. Feedback connections to the lateral geniculate nucleus and cortical response properties. Science 286, 1552–1554.

Nalatore, H., Ding, M., Rangarajan, G., 2007. Mitigating the effects of measurement noise on granger causality. Physical Review E—Statistical, Nonlinear, and Soft Matter Physics 75, 031123.

Nauhaus, I., Busse, L., Carandini, M., Ringach, D.L., 2009. Stimulus contrast modulates functional connectivity in visual cortex. Nature neuroscience 12, 70–76.

Olsen, S.R., Bortone, D.S., Adesnik, H., Scanziani, M., 2012. Gain control by layer six in cortical circuits of vision. Nature 483, 47–52.

Ozeki, H., Finn, I.M., Schaffer, E.S., Miller, K.D., Ferster, D., 2009. Inhibitory stabilization of the cortical network underlies visual surround suppression. Neuron 62, 578–592.

Pang, Z., Alamia, A., VanRullen, R., 2020. Turning the stimulus on and off changes the direction of α traveling waves. Eneuro 7.

Park, Y., Cha, Y., Kim, H., Kim, Y., Woo, J.H., Cho, H., Mashour, G.A., Xu, T., Lee, U., Hong, S.J., et al., 2025. Sub-second fluctuation between top-down and bottom-up modes distinguishes diverse human brain states. bioRxiv.

Patel, J., Fujisawa, S., Berényi, A., Royer, S., Buzsáki, G., 2012. Traveling theta waves along the entire septotemporal axis of the hippocampus. Neuron 75, 410–417.

Pesaran, B., Vinck, M., Einevoll, G.T., Sirota, A., Fries, P., Siegel, M., Truccolo, W., Schroeder, C.E., Srinivasan, R., 2018. Investigating large-scale brain dynamics using field potential recordings: analysis and interpretation. Nature neuroscience 21, 903–919.

Pfurtscheller, G., Stancak Jr, A., Neuper, C., 1996. Event-related synchronization (ers) in the alpha band—an electrophysiological correlate of cortical idling: a review. International journal of psychophysiology 24, 39–46.

Raichle, M.E., 2015. The brain’s default mode network. Annual review of neuroscience 38, 433–447.

Rao, R.P., Ballard, D.H., 1999. Predictive coding in the visual cortex: a functional interpretation of some extra-classical receptive-field effects. Nature neuroscience 2, 79–87.

Rauss, K., Schwartz, S., Pourtois, G., 2011. Top-down effects on early visual processing in humans: A predictive coding framework. Neuroscience & Biobehavioral Reviews 35, 1237–1253.

Reynaud, A., Masson, G.S., Chavane, F., 2012. Dynamics of local input normalization result from balanced short-and long-range intracortical interactions in area v1. Journal of neuroscience 32, 12558–12569.

Richardson, K.A., Schiff, S.J., Gluckman, B.J., 2005. Control of traveling waves in the mammalian cortex. Physical review letters 94, 028103.

Rubino, D., Robbins, K.A., Hatsopoulos, N.G., 2006. Propagating waves mediate information transfer in the motor cortex. Nature neuroscience 9, 1549–1557.

Schneider, M., Broggini, A.C., Dann, B., Tzanou, A., Uran, C., Sheshadri, S., Scherberger, H., Vinck, M., 2021. A mechanism for inter-areal coherence through communication based on connectivity and oscillatory power. Neuron 109, 4050–4067.

Schneider, M., Tzanou, A., Uran, C., Vinck, M., 2023. Cell-type-specific propagation of visual flicker. Cell Reports 42.

Schreiber, T., 2000. Measuring information transfer. Physical review letters 85, 461.

Schwenk, J.C., Alamia, A., 2024. A hierarchical multiscale model of forward and backward alpha-band traveling waves in the visual system. bioRxiv, 2024–11.

Schwenk, J.C., Alamia, A., 2026. Detection and quantification of planar traveling waves in the eeg using spherical phase fitting. Journal of Neuroscience Methods, 110806.

Seth, A.K., Barrett, A.B., Barnett, L., 2015. Granger causality analysis in neuroscience and neuroimaging. Journal of Neuroscience 35, 3293–3297.

Shao, Z., Burkhalter, A., 1996. Different balance of excitation and inhibition in forward and feedback circuits of rat visual cortex. Journal of Neuroscience 16, 7353–7365.

Shojaie, A., Fox, E.B., 2022. Granger causality: A review and recent advances. Annual Review of Statistics and Its Application 9, 289–319.

Solomon, E.A., Stein, J.M., Das, S., Gorniak, R., Sperling, M.R., Worrell, G., Inman, C.S., Tan, R.J., Jobst, B.C., Rizzuto, D.S., et al., 2019. Dynamic theta networks in the human medial temporal lobe support episodic memory. Current Biology 29, 1100–1111.

Spyropoulos, G., Saponati, M., Dowdall, J.R., Schölvinck, M.L., Bosman, C.A., Lima, B., Peter, A., Onorato, I., Klon-Lipok, J., Roese, R., Neuenschwander, S., Fries, P., Vinck, M., 2022. Spontaneous variability in gamma dynamics described by a damped harmonic oscillator driven by noise. Nature communications 13, 1–18.

Spyropoulos, G., Schneider, M., van Kempen, J., Gieselmann, M.A., Thiele, A., Vinck, M., 2024. Distinct feedforward and feedback pathways for cell-type specific attention effects. Neuron.

Stokes, P.A., Purdon, P.L., 2017. A study of problems encountered in granger causality analysis from a neuroscience perspective. Proceedings of the national academy of sciences 114, E7063–E7072.

Sugihara, G., May, R., Ye, H., Hsieh, C.h., Deyle, E., Fogarty, M., Munch, S., 2012. Detecting causality in complex ecosystems. science 338, 496–500.

Townsend, R.G., Gong, P., 2018. Detection and analysis of spatiotemporal patterns in brain activity. PLoS computational biology 14, e1006643.

Tsodyks, M.V., Skaggs, W.E., Sejnowski, T.J., McNaughton, B.L., 1997. Paradoxical effects of external modulation of inhibitory interneurons. Journal of neuroscience 17, 4382–4388.

Vezoli, J., Magrou, L., Wang, X.J., Knoblauch, K., Vinck, M., Kennedy, H., 2020. Cortical hierarchy and the dual counterstream architecture. bioRxiv, 2020–04.

Vezoli, J., Vinck, M., Bosman, C.A., Bastos, A.M., Lewis, C.M., Kennedy, H., Fries, P., 2021. Brain rhythms define distinct interaction networks with differential dependence on anatomy. Neuron 109, 3862–3878.

Vinck, M., Huurdeman, L., Bosman, C.A., Fries, P., Battaglia, F.P., Pennartz, C.M., Tiesinga, P.H., 2015. How to detect the granger-causal flow direction in the presence of additive noise? Neuroimage 108, 301–318.

Vinck, M., Uran, C., Spyropoulos, G., Onorato, I., Broggini, A.C., Schneider, M., Canales-Johnson, A., 2023. Principles of large-scale neural interactions. Neuron 111, 987–1002.

Voloh, B., Valiante, T.A., Everling, S., Womelsdorf, T., 2015. Theta–gamma coordination between anterior cingulate and prefrontal cortex indexes correct attention shifts. Proceedings of the National Academy of Sciences 112, 8457–8462.

Wilmes, K.A., Clopath, C., 2019. Inhibitory microcircuits for top-down plasticity of sensory representations. Nature communications 10, 5055.

Womelsdorf, T., Johnston, K., Vinck, M., Everling, S., 2010a. Theta-activity in anterior cingulate cortex predicts task rules and their adjustments following errors. Proceedings of the National Academy of Sciences 107, 5248–5253.

Womelsdorf, T., Vinck, M., Leung, L.S., Everling, S., 2010b. Selective theta-synchronization of choice-relevant information sub-serves goal-directed behavior. Frontiers in human neuroscience 4, 210.

Xu, Y., Long, X., Feng, J., Gong, P., 2023. Interacting spiral wave patterns underlie complex brain dynamics and are related to cognitive processing. Nature human behaviour 7, 1196–1215.

Ye, Z., Bull, M.S., Li, A., Birman, D., Daigle, T.L., Tasic, B., Zeng, H., Steinmetz, N.A., 2023. Brain-wide topographic coordination of traveling spiral waves. bioRxiv, 2023–12.

Zeng, Y., Sauseng, P., Alamia, A., 2024. Alpha traveling waves during working memory: Disentangling bottom-up gating and top-down gain control. Journal of Neuroscience 44.

Zhang, H., Watrous, A.J., Patel, A., Jacobs, J., 2018. Theta and alpha oscillations are traveling waves in the human neocortex. Neuron 98, 1269–1281.

Zhigalov, A., Jensen, O., 2020. Alpha oscillations do not implement gain control in early visual cortex but rather gating in parietooccipital regions. Human Brain Mapping 41, 5176–5186.

